# Long duration environmental biosensing by recording analyte detection in DNA using recombinase memory

**DOI:** 10.1101/2023.08.10.552812

**Authors:** Prashant Bharadwaj Kalvapalle, Swetha Sridhar, Jonathan J. Silberg, Lauren B. Stadler

**Affiliations:** Systems, Synthetic, and Physical Biology Graduate Program, Rice University, 6100 Main Street, MS-180, Houston, Texas 77005; Department of BioSciences, Rice University, 6100 Main Street, MS-140, Houston, TX, 77005; Department of Bioengineering, Rice University, 6100 Main Street, MS-142, Houston, TX, 77005; Department of Chemical and Biomolecular Engineering, Rice University, 6100 Main Street, MS-362, Houston, TX, 77005; Department of Civil and Environmental Engineering, Rice University, 6100 Main Street, MS-519 Houston, TX, 77005

**Author notes:** To whom correspondence should be addressed: Lauren B. Stadler and Jonathan J. Silberg.

**Keywords:** biosensor, integrase, genetic memory, quorum sensing, recombinase, wastewater, and synthetic biology

## Abstract

Microbial biosensors that convert environmental information into real-time visual outputs are limited in their sensing abilities in complex environments, such as soil and wastewater. Alternative reporter outputs are needed that stably record the presence of analytes. Here, we test the performance of recombinase-memory biosensors that sense a sugar (arabinose) and a microbial communication molecule (3-oxo-C12- homoserine lactone) over 8 days (∼70 generations) following analyte exposure. These biosensors use analyte sensing to trigger the expression of a recombinase which flips a segment of DNA, creating a genetic memory, and initiates fluorescent protein expression. The initial designs failed over time due to unintended DNA flipping in the absence of the analyte and loss of the flipped state after exposure to the analyte. Biosensor performance was improved by decreasing recombinase expression, removing the fluorescent protein output, and using qPCR to read out stored information. Application of memory biosensors in wastewater isolates achieved memory of analyte exposure in an uncharacterized *Pseudomonas* isolate. By returning these engineered isolates to their native environments, recombinase-memory systems are expected to enable longer duration and *in situ* investigation of microbial signaling, community shifts, and gene transfer beyond the reach of traditional environmental biosensors.

**IMPORTANCE:** Living microbial sensors can monitor chemicals and biomolecules in the environment in real-time, but they remain limited in their ability to function on the week, month, and year timescales. To determine if environmental microbes can be programmed to record the detection of analytes over longer timescales, we evaluated whether the sensing of a microbial signaling molecule could be recorded through a DNA rearrangement. We show that off-the-shelf DNA memory is suboptimal for long-duration information storage, use iterative design to enable robust functioning over more than a week, and demonstrate DNA memory in an uncharacterized wastewater *Pseudomonas* isolate. Memory biosensors will be useful for monitoring the role of quorum sensing in wastewater biofilm formation, and variations of this design are expected to enable studies of ecological processes *in situ* that are currently challenging to monitor using real-time biosensors and analytical instruments.

## INTRODUCTION

Bacteria continuously sense and respond to dynamic changes in their environment using cytosolic and extracellular sensing systems (1–3). Engineers have leveraged the exquisite sensing capabilities of microorganisms through synthetic biology to program them to sense and report on specific environmental conditions and analytes (4), ranging from the presence of toxic metals (5), organic pollutants (6), and antibiotics (7) to essential nutrients (8) and microbial communication signals (9). With most biosensors, reporting is achieved by coupling the sensing of a specific environmental chemical to the production of a visual output, such as enzymes that produce colored products (10,11), luminescent enzymes (12,13), and fluorescent proteins (14). Biosensors with visual outputs have enabled rapid detection of analytes within drinking water, serum, and milk with minimal pre-processing (6). Strategies have been developed to extend visual-reporting biosensors to hard-to-image materials by creating soil microcosms or rhizotrons with windows that allow imaging of microbes on the soil surface (9,15). However, visual outputs can only be used in transparent or accessible environments (12), limiting their applications in soil, wastewater, and sediment. Biosensors that record information in DNA represent an emerging technology with the potential for environmental sensing in turbid and hard-to-access environments over time scales relevant to ecological processes (Figure 1A).

**Figure 1.**
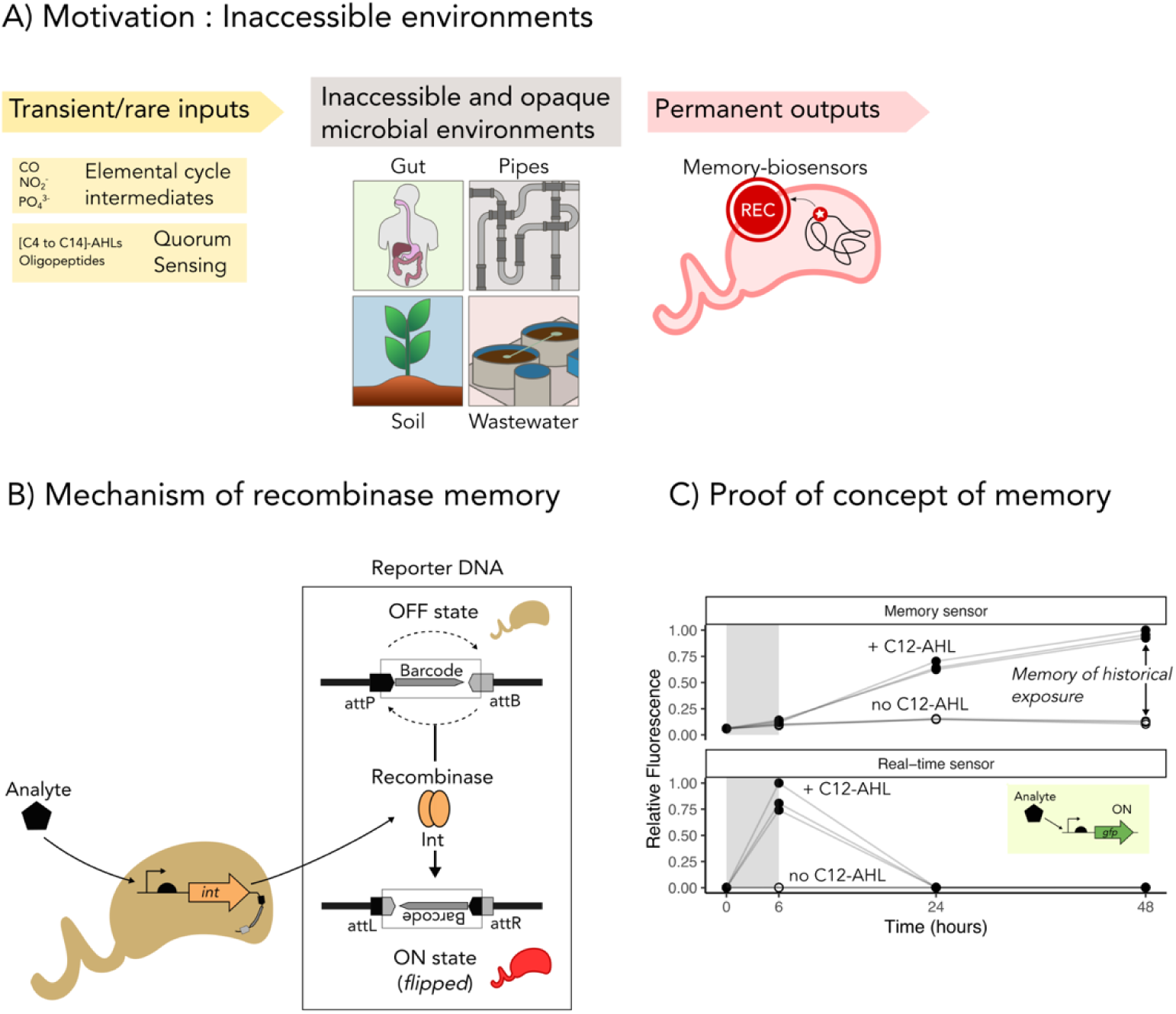
**Recombinase memory stores information of historical analyte exposure**. (**A**) Memory biosensors can be deployed into inaccessible, hard-to-image environments for undisturbed monitoring of ecological processes by coupling the detection of transient and rare chemical inputs into permanent DNA modifications. (**B**) A recombinase-memory biosensor works by conditionally expressing a recombinase enzyme when it senses an analyte. The recombinase binds the DNA attachment sites *attP and attB* and reverses the DNA segment flanked by the sites, thereby encoding information as a genetic memory in the DNA. (**C**) A memory sensor provides information on historical exposures to environmental chemicals (*top*), in contrast to a real-time reporter which only provides information during chemical exposure (*bottom*). Data is shown for reporters of the quorum sensing molecule, 3-oxo-C12-L-homoserine-lactone (C12-AHL). Sensor outputs were measured indirectly by monitoring cellular fluorescence arising from DNA flipping. Three independent cultures of both sensors were exposed to 1 µM C12-AHL for 6 hours in the exponential phase (gray), prior to washing and subculturing for 2 passages. While the memory sensor retained the fluorescence after the analyte was removed, the real-time sensor’s output was not significantly different from baseline at 48 hours (paired sample two-tailed t-test, p = 0.84).

Microbes have been engineered to record information about sensed analytes in their DNA using a wide range of enzymes, such as recombinases (16,17), CRISPR nucleases (18,19), CRISPR integrases (20,21), and polymerases (22,23). Microbes have also been engineered to record information about uptake of mobile DNA in their 16S ribosomal RNA using ribozymes (24). Among these two biological information storage approaches, information coded in DNA is more stable because the modified nucleic acid is inherited in daughter cells as the programmed cells divide. Thus, it can be retrieved after incubations of varying duration. In contrast, RNA memory is less permanent as the RNA information is degraded within cells. While DNA memory is appealing to use for environmental biosensing, since it records information about historic exposure to analytes (25), it has only been applied in a small number of environmental microbes and materials.

Recombinase-memory biosensors represent one of the earliest innovations of synthetic cellular recording (16,17). With these DNA memory devices, the conditional expression of a recombinase is used to catalyze the flipping of a DNA sequence flanked by two recognition sites (26,27). The simplicity of these systems have made them appealing for environmental applications (Figure 1B), because they only require low-level expression of a single enzyme and short ∼40-70 bp recognition sites within the DNA substrate for this enzyme (27,28); no other host derived biomolecules are needed. Recombinase memory was first applied in *Vibrio cholerae* within the mouse gut to sense iron limitation (16). Upon sensing the analyte, the recombinase activated an antibiotic resistance gene, which was detected in various regions of the intestine after sacrificing the mouse and selecting for antibiotic resistant microbes (16). Recombinase memory has also been used to identify virulence genes in pathogens (29–31), to identify symbiosis-activated genes in *Sinorhizobium meliloti* (17), to quantify root exudates (32) and contaminants (33) in soils, and to mark specific cell populations in embryos for tracing their lineage to organ formation (34,35). Although recombinase studies have achieved biosensing within different hard-to-image environments, they have largely involved short, two-day incubations. Currently, it remains unclear whether week-long applications of recombinase memory can be achieved with existing designs or if new systems must be created to achieve longer duration biosensing.

To better understand how to create memory sensors that retain information about transient analyte exposure on the week time scale (Figure 1C), we built recombinase-memory biosensors that write information in DNA about exposure to the sugar arabinose and to the signal 3-oxo-C12-homoserine lactone (C12-AHL), a quorum sensing molecule used in microbial communication. In environments such as wastewater, quorum sensing regulates many processes, including biofilm formation (36,37), flocculation (38), and pollutant biotransformation (39). We investigated the fidelity of this recombinase memory when DNA flipping was used to switch on production of a protein reporter and characterized the challenges associated with memory stability. We show that removing the burden of protein expression from the memory system enabled it to function stably for up to 8 days. We further demonstrate that the memory system can be applied in wastewater isolates and that qPCR can be used to rapidly read out the information stored. This work paves the way for the application of memory biosensors in wastewater for monitoring fundamental biological processes *in situ*, such as the signals that trigger biofilm formation, and transient toxic chemicals that are a threat to public health.

## RESULTS

### Recombinase memory records analog information

The basic unit of recombinase memory is a segment of DNA that can code binary information: OFF or ON state. In a population of bacteria, multiple digital recordings within each bacteria can result in an analog record of the analyte concentration exposure. Each bacteria contains multiple copies of the plasmid DNA (∼5 to 15), and there can be billions of bacteria per milliliter of culture. Thus, the quantification of memory depends on the fraction of the cells within the population and fraction of plasmids in each cell having DNA in the ON state. Figure 2A illustrates how transient exposure to small analyte concentrations is expected to only flip a small fraction of the bacteria within the population, while higher concentrations flip an increasingly larger fraction of bacteria into ON state, until the whole population is converted to the ON state.

**Figure 2.**
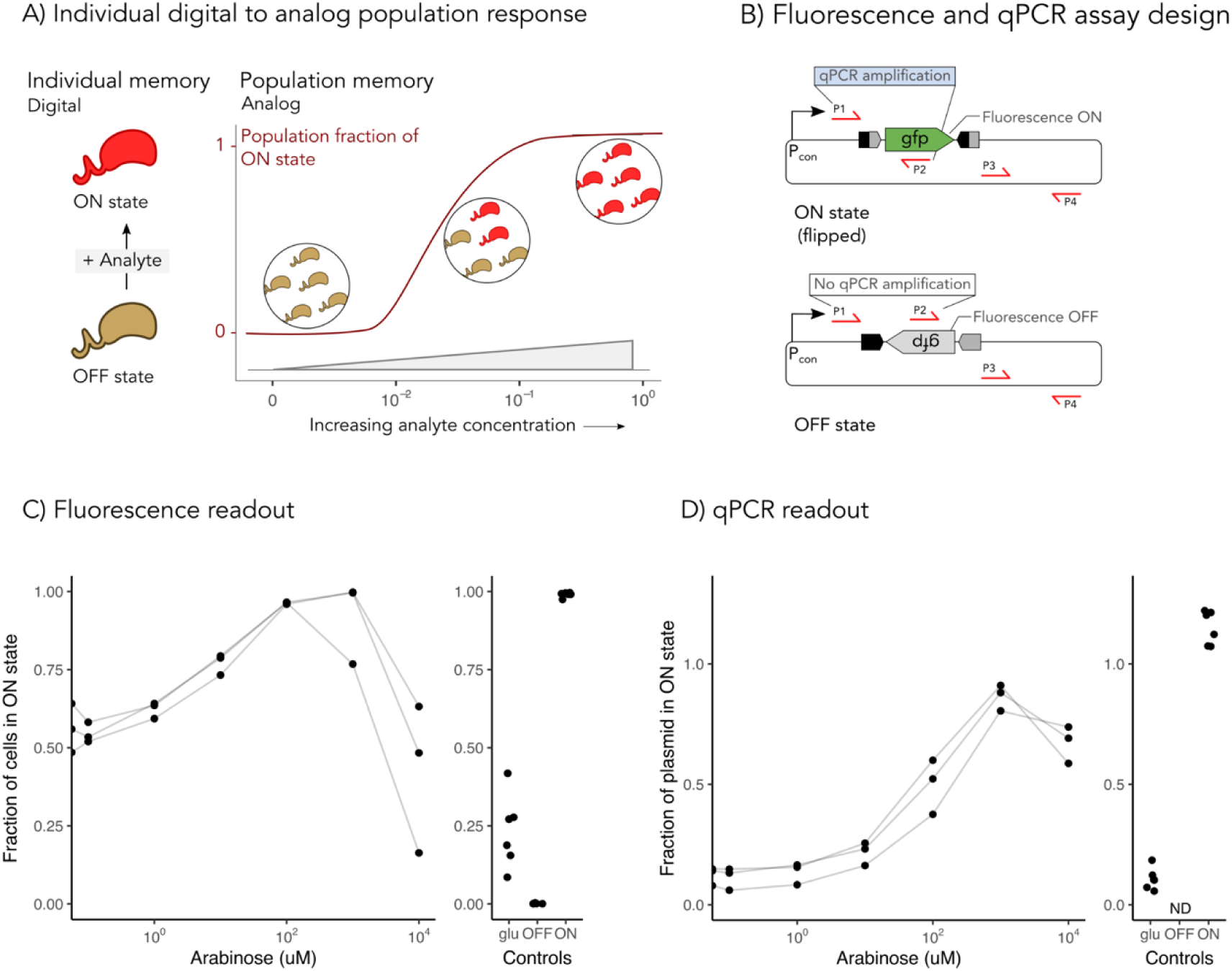
Recombinase memory stores analog information. (**A**) A digital memory storage device coded within individual bacteria can code analog information at the population level. Exposure of cells in the OFF state (brown) to low analyte concentrations marks a subset of the cells with the memory, switching them to the ON state (red), while higher concentrations increase the fraction of cells in the population to the ON state. (**B**) Primer sets were designed to read out memory using qPCR. The first primer pair anneals outside the region that is flipped to code a memory (P1) and inside the flipped DNA (P2), such that a product is only formed with the ON state. The second primer set (P3 and P4) was designed to quantify the total reporter plasmid for normalization. (**C**) The ON state was read out using fluorescence after 24 hours of exposure to a range of concentrations of the input analyte, arabinose. Flow cytometry data from 3 independent replicate cultures is shown, where the ON state is defined as fluorescence values >99 percentile of the OFF state control. The line connects data from related cultures. (**D**) The ON state fraction was quantified using qPCR as the [copies of flipped state DNA normalized to total copies of the plasmid]. Controls are shown for cells transformed with a plasmid in the ON state (ON), OFF state (OFF), or the memory biosensor with glucose (glu) for repressing the pBAD promoter. ND = non-detectable signal.

To evaluate recombinase memory as an analog sensor for environmental applications, we characterized the performance of a previously described recombinase-memory biosensor for the sugar arabinose (40), and we built another sensor for the microbial communication molecule 3-oxo-C12-homoserine lactone (C12-AHL). We characterized several memory-biosensor properties in *Escherichia coli*, including: (1) the stability of the OFF state in the absence of the analyte, (2) the sensitivity to input analyte concentration, (3) the dynamic range of the memory signal following analyte exposure, (4) the stability of the ON state following recording of analyte detection, and (5) the function of the sensor plasmid across different bacteria. To quantify the sensitivity and the dynamic properties of the biosensors, we exposed each memory biosensor to different concentrations of their respective analytes (arabinose or C12-AHL) and quantified the memory at different times after the exposure.

To compare different approaches for reading out the memory, two methods were used (Figure 2B). First, an indirect method was used in which green fluorescent protein (GFP) production was turned ON by the recombinase and monitored using the change in fluorescence of the cell. We also used a more direct method that uses quantitative PCR (qPCR) to determine the amount of DNA in each state. With the qPCR approach, we detected the ON state of the memory by designing a primer pair that binds outside and inside the flipping region, such that amplification only occurs when the memory is in the ON state and after flipping has occurred.

The arabinose sensor performance was evaluated using fluorescence and qPCR readouts. Figures S1 and S2 show the complete flow cytometry distribution and absolute qPCR quantification data, respectively. With both methods of detection, increasing concentrations of the analyte led to an increased fraction of the population in the ON state. We quantified the dynamic range of the memory as the ratio of the signal at the maximum analyte concentration (saturated) to the signal of the no analyte control (Figures 2C-2D). With the arabinose sensor, we found that both outputs presented a similar dynamic range. When the same characterization was performed with the C12-AHL sensor (Figure S3), a similar trend was observed, with increasing analyte concentrations initially showing more flipping to the ON state, indicated by higher fluorescence intensity, followed by a downward trend at the highest analyte concentrations. These results show that memory biosensors can record analog information about analyte exposure by flipping different fractions of the DNA in the population into the ON state.

### Memory of analyte exposure is unstable

To assess the stability of the OFF and ON states over time, we grew our initial *E. coli* biosensors in serial batch cultures over 8 days (Figure 3A). With this protocol, we exposed the biosensors to analyte at the beginning of the experiment for one day, and subsequently passaged them every 24 hours by diluting each stationary phase culture into fresh growth medium lacking analyte. By repeating this protocol for 8 days, we estimate that ∼72 generations of growth occurred following analyte exposure. Each day, we quantified the memory signal using single cell fluorescence measurements by flow cytometry and qPCR. With the arabinose sensor (Figure 3B, S4A), we observed a signal in the absence of analyte at the beginning of the experiment, which peaked at day 2 and decreased thereafter. When this biosensor was exposed to arabinose, a strong signal was initially observed immediately upon adding the arabinose. However, this signal decreased exponentially, with a half life of ∼1 day for all replicate experiments. After two days, the arabinose-induced signal could no longer be differentiated from the background OFF state signal. With the C12-AHL sensor (Figure 3C, S4B), we also observed a signal in the absence of the analyte, although this background signal was more stable than observed with the arabinose sensor. With this sensor, the signal following analyte exposure was more stable. The AHL-induced signal remained significantly higher than the OFF state signal until day 6. It decreased by 50% between days 5 and 8 for the different replicate experiments. These results show that both the OFF and ON states of the memory biosensor are unstable over week-long incubations, illustrating the limitations of these systems.

**Figure 3.**
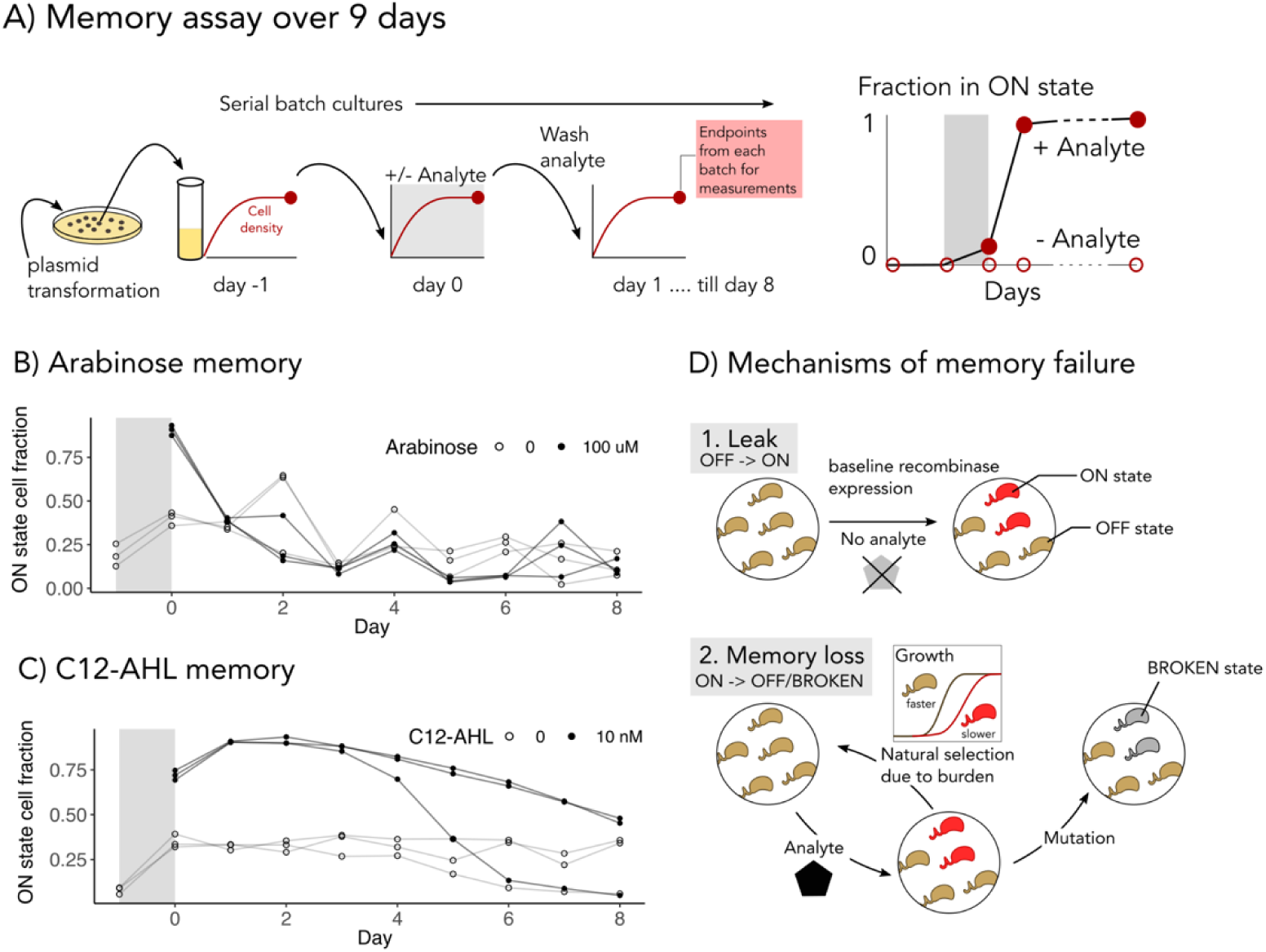
Week-long incubations led to memory loss. (**A**) Experimental design for testing memory stability. Biosensor were grown for ∼72 generations using serial batch cultures, with the analyte present only on the first day (shaded region). (**B**) The arabinose-memory sensor was tested using three independent replicate cultures following one day of exposure to 100 uM arabinose (filled circles) or no arabinose (open circles). Samples from the same biological replicates are connected by a line. The fraction of ON state cells expressing GFP were read-out by flow cytometry at the end of each day (Figure S4). The ON state was unstable, since it only remained significantly higher than no arabinose levels on days 0 and 1 (paired sample one-tailed t-test, p < 0.03). The OFF state was also unstable, as the uninduced cells presented fluorescence. (**C**) The C12-AHL memory sensor was characterized using 10 nM analyte. Both the ON and OFF states were unstable. However, the memory loss was slower, with significant difference between AHL exposed and unexposed samples until day 6 (paired sample one-tailed t-test, p < 0.03). (**D**) Mechanisms predicted to underlie memory instability. (1) Leaky recombinase expression in the absence of the analyte is predicted to yield a signal in the OFF state (brown). With time, the recombinase accumulates to sufficient levels to flip the memory to ON state (red). (2) The metabolic burden of fluorescent protein expression is predicted to make the ON state unstable. Cells in the ON state are either outcompeted by cells in the OFF state, or they accumulate mutations that abolish fluorescent protein expression resulting in a BROKEN state (gray).

Our findings implicate different mechanisms as responsible for the instability of the OFF and ON states (Figure 3D). The OFF state is thought to be unstable because there is sufficient baseline expression of the recombinase in the absence of the analyte to flip some of the DNA. The ON state is thought to be unstable due to the fitness burden that arises when expressing a fluorescent protein, which results in selection pressure favoring faster growing OFF state cells. Also, the fitness burden was thought to induce mutations that abolished the fluorescence of cells that had been switched to the ON state, termed as BROKEN states. After exposure to analyte, we observed a large deletion of ∼1.4 kb in the memory reporter plasmid on day 8 (Figure S5), supporting the concept of a BROKEN state. Taken together, our results suggest that long-duration memory systems are challenging to implement due to challenges with both the OFF and ON states.

### Memory is more stable when the protein output is removed

To learn how to improve the stability of the recorded information, we created alternative designs for the memory biosensor. We had three goals with these designs: (1) decrease the baseline recombinase expression, (2) minimize sequence repeats and remove non-functional intergenic DNA to decrease chances of recombination that lead to the BROKEN state over time, and (3) consolidate the sensor system into a single mobilizable plasmid with a broad-host range origin of replication (pBBR1) to enable its portability into environmental isolates.

To investigate how decreasing the background recombinase expression affects memory performance, we mutated the recombinase’s ribosome binding site (RBS). A library of 56 RBS variants was designed that contained variants predicted to present decreased translation initiation rates (41). Among these, five presented a decrease in the background GFP signal in the absence of analyte (Figure S6A). These refined systems presented a ≥10-fold increase in GFP signal upon analyte addition (Figure S6B), all of which had gains in signal that were greater than the original system tested. When these new designs were used to evaluate the stability of the memory system over ten days, we found that they uniformly presented a lower background GFP signal (Figure S7). These designs recorded detection of an analyte like the original design, and they presented transient information storage which rapidly decayed over two days. One of the variants was able to record the detection of two sequential pulses of analyte; the parental design could not accomplish this type of information storage. These results demonstrate that decreasing the translation initiation of the recombinase is an effective strategy to decrease the background signal (*i.e.*, improve OFF state stability). However, this change alone did not extend the stability of the stored information due to ON state instability.

To improve ON state stability, we built three new plasmids that avoided DNA sequence repeats and included only DNA that was critical to biosensor function using a hierarchical golden gate cloning method that combined each functional element, including the promoters, translation initiation sequences, coding sequence of the genes, terminators, antibiotic selection genes, and broad host range origin of replications (42). When designing these plasmids, we minimized sequence repeats with the potential to cause mutations that lead to failure of the genetic circuit (43). Figure 4A shows the three different memory designs that were built, which differ in their potential cellular burden on the host. In the first design, called “Fluorescent”, the recombinase flips a promoter, thereby turning on the production of a red fluorescent protein, mcherry2. In the second design, called “Silent”, the recombinase flips a short, noncoding DNA sequence. In the third design, called “Frugal”, the recombinase deletes the LasR analyte sensor and the recombinase, thereby creating a smaller plasmid (6.8 kb to 4.2 kb) that is expected to create a smaller resource burden on cells.

**Figure 4.**
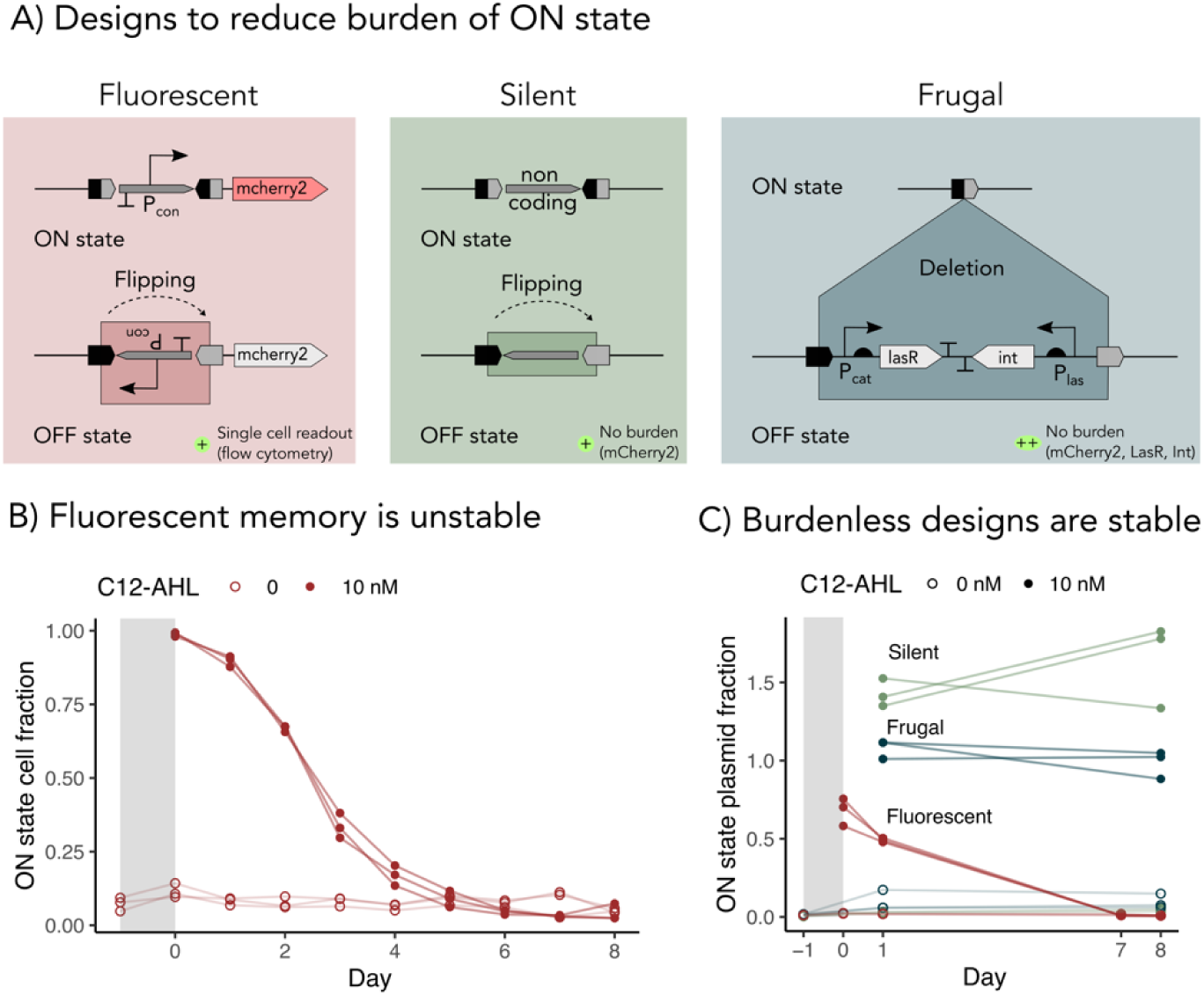
Memory designs can extend the duration of memory utility. (**A**) The Fluorescent (red), Silent (green), and Frugal (blue) biosensor designs all use a broad host range origin to enable studies in diverse bacterial species, and they lack repeat sequences that can lead to deleterious recombination. Upon sensing the analyte, the Fluorescent design produces fluorescence similar to the parent design, the Silent design flips a non-functional DNA region, and the Frugal design deletes the sensor components (*lasR* and *int* recombinase) to yield a smaller plasmid that is expected to minimize cellular burden. (**B**) The Fluorescent memory was tested for stability using three independent replicate cultures following a one day of exposure to 10 nM C12- AHL (filled circles) or no C12-AHL (open circles). The y-axis shows the fraction of cells in the ON state, gated by fluorescence (Figure S8), over the duration of the experiment. (**C**) For each design, the memory output is shown for the same experiment, measured by qPCR. The fraction of the plasmids in the ON state is defined as the copies of flipped state DNA normalized to the total copies of the plasmid; the latter was measured at the start and end of the experiment. To evaluate stability, we assessed whether the signal was significantly different when comparing days 1 and 8 of the experiment (paired sample one sided t-test on C12-AHL exposed samples for d1 > d8: Fluorescent: p = 1e-4, Silent: p = 0.8, Frugal: p = 0.16).

After sequence verifying the plasmids encoding the new designs, we tested their ability to remember exposure to an analyte (C12-AHL) over 8 days using *E. coli* and measured the memory performance by flow cytometry and qPCR. The ON state signal from cells containing the Fluorescent design rapidly decayed over time (Figure 4B), as observed in the initial designs. In contrast, we found that the ON states of the Silent and Frugal designs were both stable for 8 days following exposure to analyte, presenting signals on day 1 and 8 that differed by <30% or 0.2 fold (Figure 4C). Analysis of the fluorescence of individual cells harboring the Fluorescent design revealed low recombinase leak in the absence of analyte (Figure S8). Taken together, these results show that the Silent and Frugal designs present improved performance. This trend is thought to arise because these designs eliminated the cellular burden of fluorescent protein expression.

### Memory in wastewater environmental isolates

To enable biosensing within wastewater, we tested the optimized memory-biosensor plasmids in bacteria isolated from untreated wastewater. To obtain microbes for this test, influent wastewater was streaked onto nutrient-agar plates, and the antibiotic sensitivities of the individual isolates was characterized to identify appropriate concentrations for plasmid selections. The memory sensors were then conjugated into each isolate by mating using *E. coli* as a donor strain (Figure 5A). Once the transformed microbes were obtained on selective plates, we evaluated the ability of three wastewater isolates to stably record exposure to the analyte C12-AHL over 8 days (Figure S9). For these experiments, we evaluated information stored by the sensors using qPCR. We also compared the performance of one wastewater isolate with *E. coli* with the Fluorescent design. We found that the wastewater isolate was more stable in the ON state following exposure to analyte as compared with *E. coli* (Figure 5B, left panel). With the wastewater isolate, the half life of the information stored was ∼4 days, versus ∼1 day for *E. coli*. For the Silent design, the signal from the wastewater isolate exposed to analyte was similar to cells grown in the absence of analyte (Figure 5B, middle panel). This suggests that the cells were already in the ON state when the experiment began. This can be contrasted with the *E. coli* biosensors containing this design, which did not present a signal in the absence of analyte. With the Frugal design, the signal in the wastewater isolate increased upon addition of analyte (Figure 5B, right panel). However, this design presented a signal that increased with time in the absence of analyte as well. In contrast, the *E. coli* sensors presented a memory signal that remained higher than the background signal after eight days. These results show that two of our three memory biosensor designs yield a signal within wastewater isolates without optimization.

**Figure 5.**
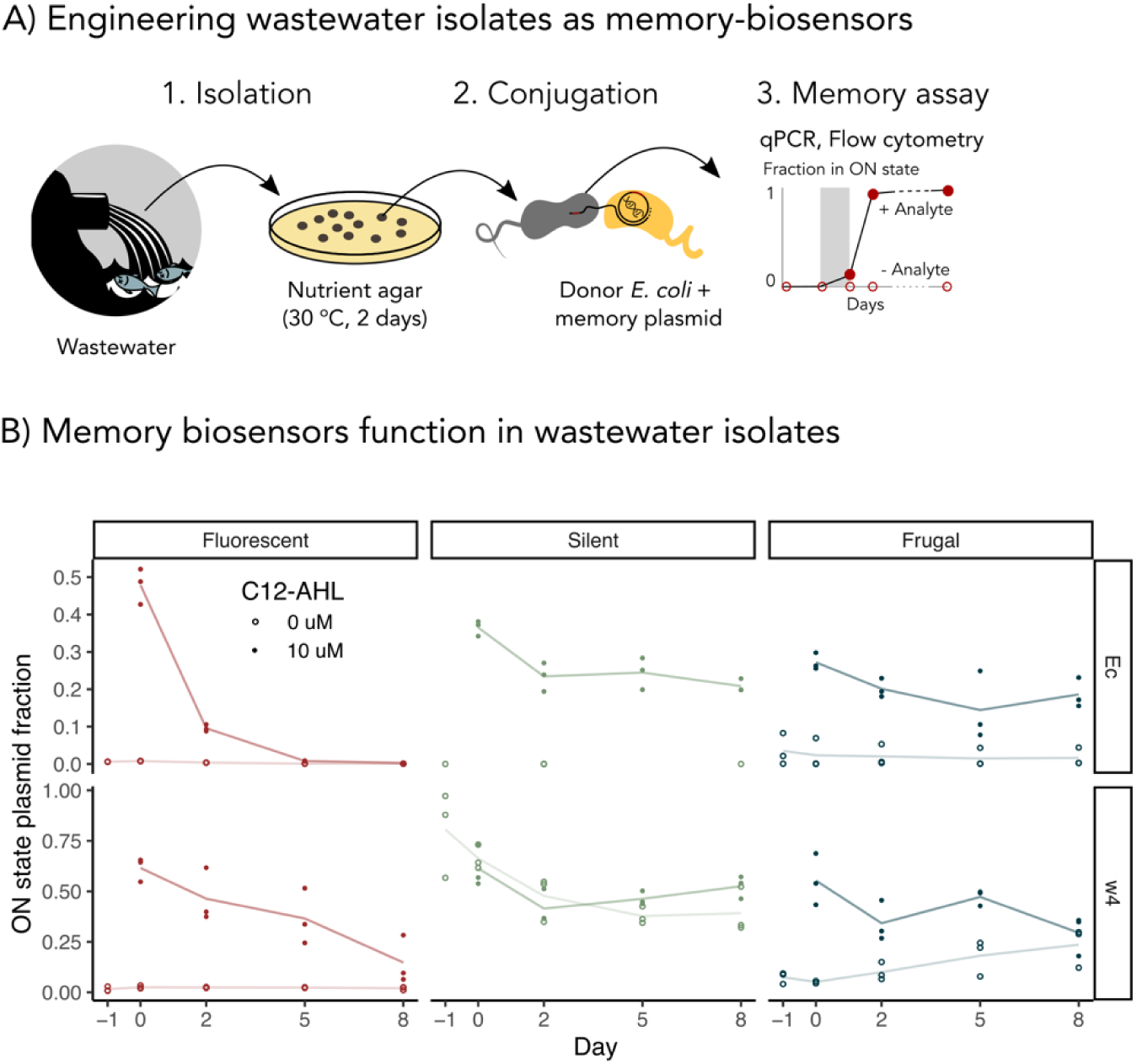
Memory designs function in wastewater microbes. (**A**) To test memory in wastewater isolates: (1) wastewater was streaked on nutrient agar plates to isolate colonies and characterized using16S rRNA sequencing, (2) the memory-biosensor plasmids were conjugated from *E. coli* donors into the wastewater isolates, and (3) memory performance was characterized using qPCR. (**B**) A comparison of the memory designs in *E. coli* (top) and one wastewater isolate w4 (bottom). The performance of the Fluorescent (red), Silent (green), and Frugal (blue) designs are compared. For each experiment, we measured the fraction of plasmid in the ON state as in Figure 4, with data from three independent cultures shown as points, and a line connecting the averages. With the Fluorescent design, analyte exposure yielded a signal that decayed with a half life of 1.0 ± 0.1 days with *E. coli* and 3.6 ± 0.8 days with the wastewater isolate (fit to a single exponential model). With the Silent design, analyte exposure yielded a signal with *E. coli* that was significantly higher than uninduced (p = 0.0044 on day 2, control not detected other days) but not for the isolate (p > 0.64). With the Frugal design, analyte exposure yielded a signal that was significantly higher than the control on all days with *E. coli* (p < 0.04) and with the isolate (p < 0.033). Isolate without exposure had a significantly higher signal on days 5 and 8, than day -1 (p < 0.048). All the p-values were obtained using a paired sample one-tailed t-test.

We hypothesized that the background signal observed with the Frugal design in the wastewater isolate could arise through two mechanisms. The genetic program could express the recombinase in the absence of the analyte, and/or the wastewater isolates could synthesize the C12-AHL analyte being sensed. To address the latter possibility, we investigated whether the different wastewater isolates produce C12-AHL by incubating the spent media from cultures of each isolate with an *E. coli* C12-AHL biosensor. With this analysis (Figure S10), the spent medium from each culture lacked sufficient C12-AHL to activate the *E. coli* fluorescence sensor, which uses the same transcriptional regulator as our memory biosensors for sensing the analyte. This finding suggests that it is more likely that the wastewater isolates with the Frugal and Silent systems present low level recombinase expression in the absence of analyte. Thus, the regulatory regions which were optimized for *E. coli* expression could be further optimized in the wastewater isolate to improve performance.

## DISCUSSION

Our results show that recombinase-memory biosensors can be engineered to record exposure to an environmentally-relevant analyte within DNA using both *E. coli* and a wastewater isolate. We also show how this form of memory for environmental analytes can be programmed to function stably for up to 8 days. Prior studies using recombinase memory tested performance over shorter day-long incubations (16,17,29,33). Here, we extended the utility of this type of memory to longer timescales by minimizing the cellular burden of the biosensor components and tightly regulating the conditional expression of the recombinase, which writes information in DNA (40). We target an environmentally-relevant analyte, the microbial communication molecule C12-AHL, which enables coordinated microbial action such as biofilm formation (38), and flocculation, which are important to wastewater treatment performance and process management (38). While real-time biosensors that generate a visual reporter have been widely used for AHL sensing, the memory biosensor described here is expected to overcome current limitations with those biosensors (9,44,45), which require continuous monitoring to detect transient analytes. In environmental materials, it is often not possible to obtain time-resolved information with real-time visual outputs due to their opaque nature. Gas-reporter outputs have been reported that overcome this challenge (46). However, they require complex analytic instrumentation to monitor biosensor outputs, whose limits of detection can constrain the information obtained from biosensors (47). In contrast, the biosensors developed herein are expected to overcome these shortcomings and enable monitoring of the wastewater processes *in situ* and over longer timescales.

Our results identify design strategies that can be used to improve the stability of recombinase-memory biosensors. First, baseline expression of the recombinase can be tuned down by decreasing the strength of translation initiation to minimize false recording that occurs in the absence of exposure to the analyte. This tuning, which can be accomplished using a thermodynamic model for translation initiation (41,48), is especially important in long duration applications of memory biosensors, as even low level recombinase expression can lead to production of significant ON state over week-long durations. Another advantage of low recombinase expression is that it avoids the fitness burden on cells when exposed to high concentrations of the analyte, which resulted in lower than expected ON state with our original designs. Second, streamlined designs that remove intergenic regions and minimize sequence repeats should be created to minimize mutations that can arise from plasmid recombination (43). Third, the cellular burden of storing information in cells should be minimized. We showed that this burden can be minimized via two different approaches. One can create a Silent design that records information in DNA without expressing a fluorescent reporter. Alternatively, one can create a Frugal design by removing the sensing components themselves in the process of recording. Among these two approaches, the Frugal design exhibited the best functioning within a wastewater isolate. These approaches were successful at improving the longevity of memory biosensors within *E. coli*, and they are expected to be generalizable for optimizing performance in other bacterial hosts.

This study represents the first time that recombinase-memory sensors have been tested in non-model bacteria, and specifically in natural isolates from wastewater. Our findings highlight the opportunities and challenges of using synthetic biology to program memory biosensing in natural isolates. While we demonstrated the application of our optimized memory biosensor in an undomesticated bacteria isolated from wastewater and showed that it performed better than our prototype *E. coli* sensors on a week-long time scale, the wastewater isolate did not perform as well as the optimized system in *E. coli*. This finding illustrates how differences in transcriptional and translational regulation across microbes from the same phylum can lead to variation in the performance of a genetic program. In the future, the memory biosensor can be refined to present greater stability in wastewater isolates and applied in real communities and materials like sludge to evaluate what individual microbes perceive in their local environments. The use of modified environmental isolates may improve the stability of the memory biosensor, as they are likely able to survive in their native environment without the challenges that non-native microbes such as a domesticated *E. coli* would encounter in such environments. In the gut environment, synthetic genetic circuit performance has been improved by targeting native microbes (49,50).

Further improvements could be explored to minimize the signal in the absence of analyte and improve the stability of the recorded information beyond what we examined. Two possible mechanisms that could trigger the background signal that were not explored in this work are: (i) the presence of native recombinases in the wastewater isolates that may cause a low level of recording, and (ii) the use of inducible promoter design with inherent baseline activity. For the former, bioinformatic analysis can be used to identify native recombinases in strains targeted for genetic programming, and *in vitro* and *in vivo* testing can be used to parse out mechanisms of native recombination that interferes with the stability of the synthetic recombinase circuits. For the latter, a non-zero baseline expression of the recombinase is inevitable with the current inducible promoter design due to the kinetics of transcription factor binding (51). This challenge can be overcome by incorporating a secondary control on recombinase expression, such as constitutive expression of an inhibitory antisense RNA that inhibits the translation of the recombinase and imposes a threshold of analyte sensing before recombinase protein is synthesized and recording occurs (51).

In the future, recombinase-memory systems that enable longer duration experiments in environmentally-representative systems can be directly applied to study microbial interactions such as cell-cell signaling that underlies the formation of biofilms widespread in drinking water and wastewater systems. By recording information using different recombinases (40,52), wastewater biosensors could be created that record information about exposure to different types of quorum sensing molecules simultaneously in the same community. The use of qPCR for reading out the information stored, which is widely used for environmental microbiology (53), is cheaper per run compared to flow cytometry analysis of fluorescent protein reporters. Thus, this approach for memory biosensor readout is expected to be more widely accessible than memory biosensors that produce visual outputs. Memory biosensors are also expected to enable fundamental and minimally-invasive monitoring of a wide range of environmental processes that are hard to study *in situ* such as horizontal gene transfer, plant-microbe interactions, and quorum sensing.

## MATERIALS AND METHODS

### Chemicals

Reagents for growth media, antibiotics, and inducers were purchased from Sigma Aldrich. Enzymes for molecular biology (BsaI-HF v2, T4 DNA ligase, Taq ligase, T5 exonuclease, and Phusion DNA polymerase) were from New England Biolabs, and qPCR master mixes were from PCR Biosystems. Primers were purchased from Sigma Aldrich, and probes were from Integrated DNA Technologies and LGC Biosearch Technologies. DNA extractions were performed using Qiagen Miniprep Kit.

### Isolating wastewater bacteria and conjugation

Untreated wastewater samples stored at 4°C were spread onto YPS-agar (4g/L yeast extract, 2g/L peptone, 25 g/L sea salts) or LB-agar medium. They were grown at room temperature (∼25°C) for 2 days, and colonies with different morphologies were re-streaked before inoculation in super optimal broth with catabolite repression (SOC) medium (54). The sensitivity of individual isolates to kanamycin and chloramphenicol were characterized by serial-dilution spotting on LB-agar plates with different concentrations of the antibiotics. Memory plasmids were transformed into the donor strain, *E. coli* MFDpir by heat shock (55), and this strain was used as a donor for conjugation. Donors and recipients were grown separately to stationary phase at 37°C and room temperature, washed in sterile-filtered phosphate buffered saline (PBS) twice (56), mixed at a 1:1 ratio, spotted onto LB-agar medium in a 96-well deep-well block, and incubated for 24 hours at room temperature. Microbial mixtures were resuspended in PBS (500 µL), washed, and spotted onto selective medium. Selected colonies were re-streaked on LB-agar plates, individual colonies were used to inoculate liquid cultures, and cultures were screened for fluorescence with C12-AHL (10 µM).

### Bacterial growth

DNA construction and assembly was performed using *E. coli* DH10B grown in LB medium. Memory experiments were performed in *E. coli* MG1655, a strain that has been maintained in labs with minimal genetic manipulation (57). For memory experiments, *E. coli* were grown in M9 minimal medium (M9-glucose) containing 0.4% w/v glucose, 0.2% casamino acids, and antibiotics at 37°C (58,59). Memory experiments in wastewater isolates were performed in LB medium incubated at 30°C. For all strains, we used 50 µg/mL kanamycin. However, the chloramphenicol concentrations varied by strain: *E. coli* (34 µg/mL), w4 isolate (100 µg/mL) and the w17 and w23 isolates (200 µg/mL). For the arabinose titrations, 0.4% glycerol was substituted for glucose in the M9 medium, since glucose represses the pBAD promoter induced by arabinose (60). To induce maximal recombinase expression in *E. coli*, 100 uM arabinose or 10 nM C12-AHL were included in the growth medium during subculture. For the wastewater biosensors, 10 uM C12-AHL was used for maximal induction. All assays were performed using 3 independent colonies (biological replicates) grown until stationary phase (∼18 to 24 hours) at 37°C in M9-glucose (500 µL) with necessary antibiotics in 96-well deep-well blocks (Analytical Sales and Services, SKU: 59623-23) that were shaking at 650 rpm.

### Plasmid design

All vectors used in this study (Table S1) were constructed using Golden Gate Assembly (42). Vectors for arabinose memory (pAra) and the GFP reporter of recombinase activity (pRec1-OFF) were obtained from Addgene (40). In pAra, the serine recombinase from *Staphylococcus haemolyticus* (Uniprot: Q4L3S2_STAHJ) is expressed under the pBAD promoter. The pBAD promoter is induced by arabinose, which binds to the transcriptional activator AraC, expressed constitutively from the same plasmid. In the pRec1-OFF plasmid, a constitutive promoter is situated upstream of the *gfp* gene, but the gene is inverted in direction with respect to the promoter, so GFP is not expressed. To create the flipped pRec1-ON plasmid as a positive control, *E. coli* DH10B was transformed with pAra and pRec1-OFF, grown in LB medium with 1 mM arabinose for 16 hours at 37°C, and streaked onto an LB-agar plates containing chloramphenicol. Individual colonies were used to inoculate LB liquid cultures containing chloramphenicol, the plasmid was extracted from individual colonies, those plasmids were transformed into DH10B to isolate pRec1-ON, and the purified plasmid was sequence verified. pAHL1 contains the same recombinase as pAra, but it is controlled by a synthetic pLas promoter from *Pseudomonas aeruginosa* (BBa_K649000, IGEM registry) (61). The pLas promoter is activated by C12-AHL when bound to LasR (BBa_C0179, IGEM registry), which is expressed constitutively from the same plasmid. The plasmids (pAHL1_M1 to pAHL1_M5) with weaker RBS were screened from the RBS library built using Gibson assembly of the amplified pAHL1 backbone without the RBS (62), with a bridging ssDNA oligo containing the RBS library (Table S2). The optimized plasmid designs (pAHL2_Fluorescent, pAHL2_Silent and pAHL2_Frugal) were assembled using hierarchical Golden Gate Assembly (42) by incorporating dsDNA fragments (Twist Biosciences) containing the *att* sites of the flipping register along with the other gene constructs: pCat-catRBS-LasR and pLas-int8. The Fluorescent version included a constitutive promoter (attB-invertible[<J23110 >tVoigtS14]-attP) next to mcherry2, the Silent version included the non coding sequences (attB-non-coding-attP), and the Frugal version had attB and *inverted*-attP sites flanking the other gene cassettes.

### Short-duration memory experiments

Real-time AHL reporter (61) and memory reporter plasmids (pAra + pRec1-OFF) were transformed into *E. coli* MG1655. Three independent colonies were inoculated into LB medium, grown until stationary phase, diluted 1:100 into LB containing 1 µM C12-AHL, and grown for 6 hours. Subsequently cultures were centrifuged, washed in PBS and subcultured twice for 18-24 hours each without the C12-AHL. Green fluorescence of all cultures resuspended in PBS at the end of each timepoint was quantified with a plate reader (Tecan Spark, excitation 488 nm, emission 511 nm, gain 140, z-position 20,000 µm). Autofluorescence background calculated from MG1655 cells was subtracted, and fluorescence was normalized to optical density (OD600) to account for differences in cell growth.

### Recording information by varying arabinose concentrations

Cells co-transformed with the conditional-recombinase (pAra) and the memory-reporter (pRec1-OFF) plasmids were grown on LB-agar plates containing glucose, and three colonies from each plate were inoculated and grown for 24 hours while shaking at 650 rpm. Cells transformed with memory-reporter plasmids in the OFF (pRec1-OFF) and ON (pRec1-ON) states were also grown as controls. These cultures were then diluted 1:100 into M9 glycerol containing arabinose at different concentrations and grown for 24 hours. Aliquots of the induced cultures were diluted 1:100 in PBS with 0.1% Tween-20 (PBST) and used for flow cytometry. The same cultures were also subjected to qPCR after resuspending in water, performing a 5x dilution, and using heat lysis.

### Flow cytometry analysis of whole cell fluorescence

Cultures in stationary phase from experiments stored at 4°C for 3-9 days were resuspended by pipetting and diluted 1:100 into 500 µL PBST in 96 well deep-well blocks. Samples were run on a spectral flow cytometer with autosampler (Sony SA3800) using the low sample pressure setting. Fluorescence wavelengths were calibrated using *E. coli* MG1655 expressing *gfpmut3*, *mcherry2*, or no fluorescent protein. To calibrate fluorescence, 8-peak beads (Spherotech, RCP-30-5A) were used to establish how fluorescence relates to standardized units of molecules of equivalent fluorophores (MEFL) using FlowCal software (63). Fluorescence data was processed using custom R scripts incorporating flowworkspace (64), openCyto (65) and ggcyto packages (66). Median values extracted from a pool of 3 biological replicates were plotted. Code used for processing is available on Github (https://github.com/ppreshant/flow_cytometry/).

### Lysis and quantitative PCR

Aliquots of M9 cultures (20 µL) were centrifuged, resuspended in water, diluted into 100 µL water in PCR plates, and heated at 95°C for 10 minutes to achieve a quick lysis and high throughput DNA extraction with small sample volumes (67). For Figure 5B, cultures in LB (20 µL) were directly diluted into water for lysis without prior resuspension. These dilute microbial lysates were used directly in the qPCR reactions. Dilute lysate (4 µL) was assayed in a 10 µl qPCR reaction with qPCRBIO Probe Mix Lo-ROX master mix (PCR Biosystems, PB20.21-01) containing 0.4 µM each primer, 0.2 uM probe, and 50 nM ROX reference dye in a Quantstudio 3 thermocycler using 40 cycles of two step PCR with 95°C for lysis and 65°C for the annealing and extension temperature. Primers flanking the att regions were designed using Primer3 (68). Three sets of primer pairs and probes were designed to amplify the ON state (flipping boundary), total copies of the reporter plasmid (plasmid ori) and *E. coli* chromosome (*dcp* gene) in a single triplex reaction, and absolute copies were obtained by fitting Cq values to a standard curve (Figure S11). The fraction of ON state plasmid was obtained by dividing the copies of the ON state by the total plasmid. Details of the qPCR primer and probe sequences are available in Table S3.

### Analysis of memory stability

To test the temporal stability of the memory, *E. coli* MG1655 was transformed with the memory reporter plasmid sets, and three independent colonies were picked and grown in M9-glucose with antibiotics (500 µL) in deep-well blocks while shaking at 650 rpm at 37°C. Every 24 hours, the cultures were diluted ∼1:500 into fresh media using 96-well replicator pins, and the process was repeated for 8 days. Arabinose or C12-AHL were included in the media only between day -1 and day 0, and cells were washed with PBS before subculturing. Aliquots of the cultures at the end of each day were diluted 1:100 in PBST and used for flow cytometry. The same cultures were also subjected to qPCR after heat-lysis.

### Statistics

Data points presented represent three biological replicates derived from independent colonies with each replicate paired between exposure vs no exposure conditions and over the time course. P-values were obtained using one or two-tailed Paired samples t-tests. In cases where week-long incubations were performed, we evaluated the significance of values observed on days minus one or one and eight.

### Software

DNA sequences were stored, annotated, aligned and assembled in-silico using Benchling cloud software. Data analysis and plotting was performed in R software using ggplot2 and Rmarkdown. Automated R based workflows were used for analysis of flow-cytometry (10.6084/m9.figshare.23681499) and qPCR data (10.6084/m9.figshare.23681496). Illustrations were prepared in Inkscape.

### Data availability

Data underlying the figures along with the processing R scripts are available upon request.

## ACKNOWLEDGEMENTS

We are grateful for support from the National Science Foundation under grants 1805901 (to LBS and JJS), 2223678 (to LBS and JJS), and 2237052 (LBS). This research was also supported by a Johnson & Johnson WiSTEM2D Award (LBS) and the Gordon and Betty Moore Foundation (JJS). Additionally, we wish to acknowledge insightful discussions from Charis Giannitsis, and plasmid design from Shyam Bhakta.

## SUPPORTING INFORMATION

## SUPPLEMENTAL MATERIALS LIST

**Figure S1.**
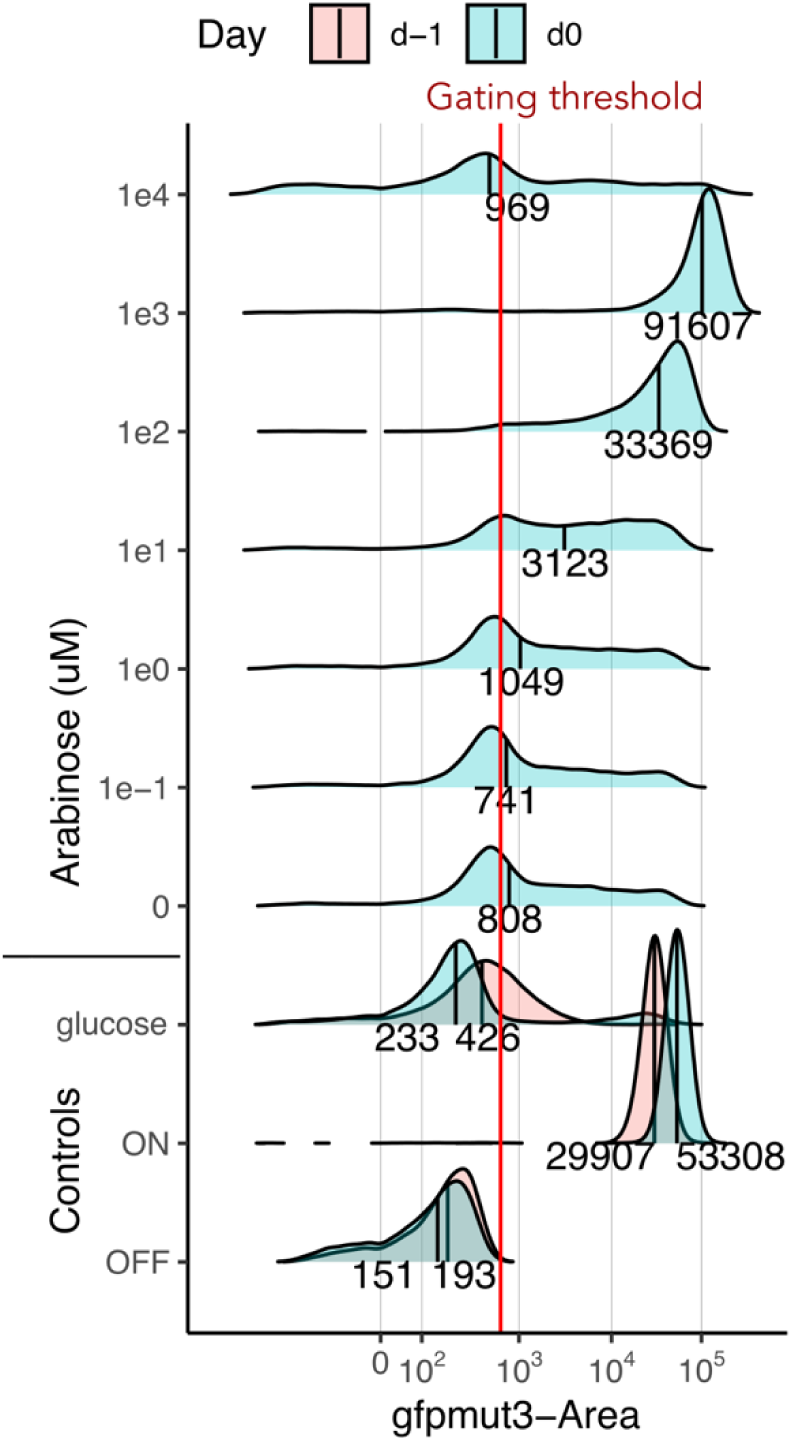
Flow cytometry distribution of arabinose memory. Ridgeline plots show the distribution of fluorescence in the arabinose-memory biosensor cell populations (pAra + pRec1-OFF) when exposed to different arabinose concentrations. The x-axis shows the fluorescence intensity, while the y-axis shows the relative abundances of cells at each intensity. The relative cell abundances of each sample were independently normalized, such that peak amplitudes are not comparable across different experiments. The vertical line within each peak and the numbers below each peak indicate the median intensity values. Three controls are shown, including ON which is pRec1-ON, the OFF which is pRec1-OFF, and glucose which is the arabinose-memory biosensor in the presence of 0.5% glucose, which represses expression. The signals from controls were measured for two days plotted and are shown in pink (day -1, d-1) and blue (day 0, d0). Cells exposed to higher arabinose concentrations shift to higher fluorescence, except with the highest concentration (1 mM arabinose); this trend is thought to arise due to the burden of recombinase expression at the highest analyte concentrations. We gated all distributions to 99 percentile of the d-1, OFF state control.

**Figure S2.**
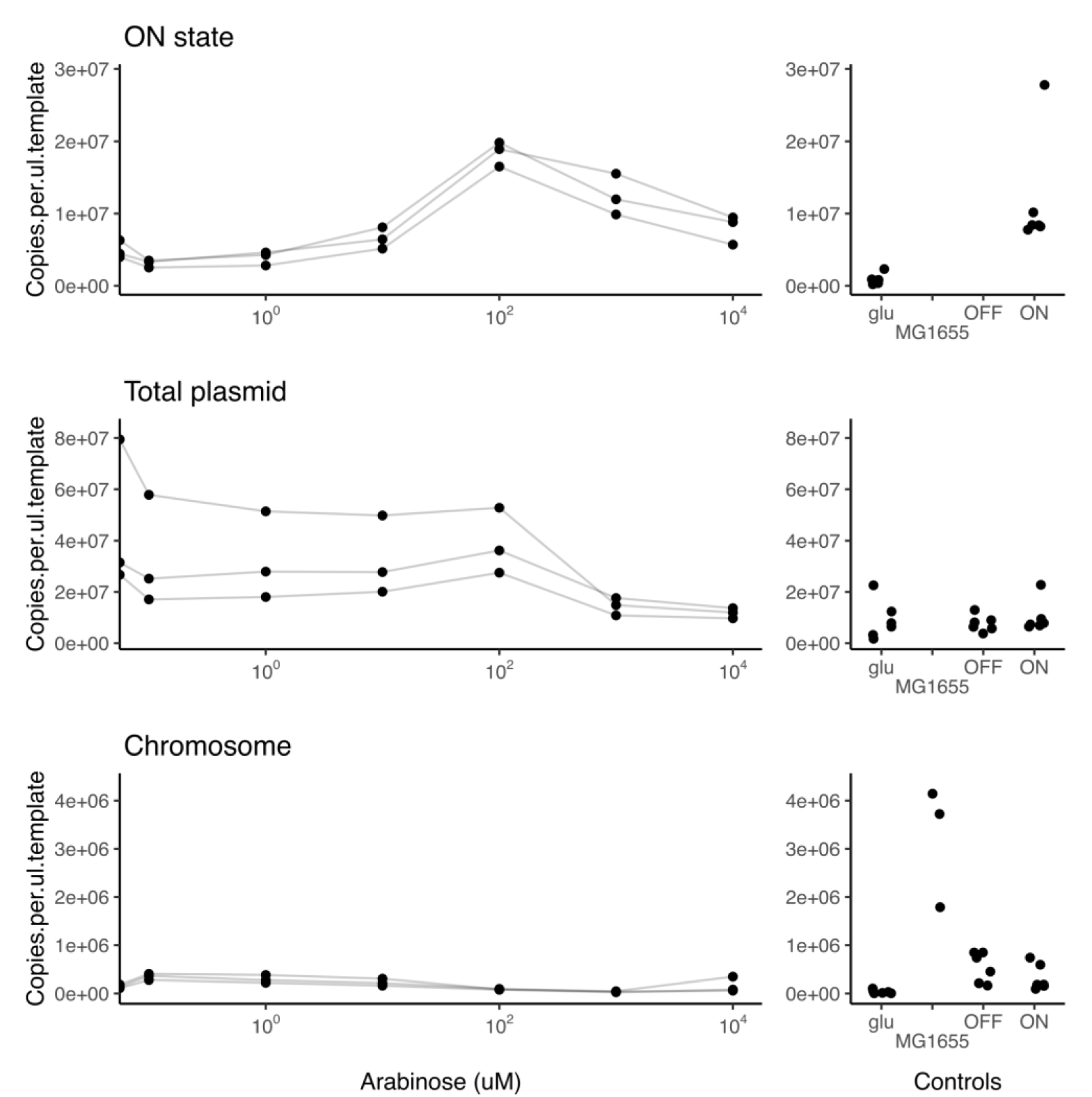
Copy number data of arabinose analog memory. We measured the copy numbers of plasmid or chromosomal targets within the arabinose-memory cells using a multiplexed qPCR assay that employed the primer pairs shown in Table S3 (ON state = Flipped-v0, total plasmid = Backbone-v0, and chromosome = Chromosome-v0). Arabinose-induced samples are shown in the left panels, while controls are shown in the right panels. The ON state and total plasmid copies increased until 100 uM arabinose and decreased at higher concentrations. This latter trend is interpreted as arising from the burden of the integrase expression at the highest analyte concentrations. Copy numbers of the chromosome were highest in empty MG1655 cells and were lower in all plasmid-carrying cells. When points are not shown, this indicates no signal using qPCR.

**Figure S3.**
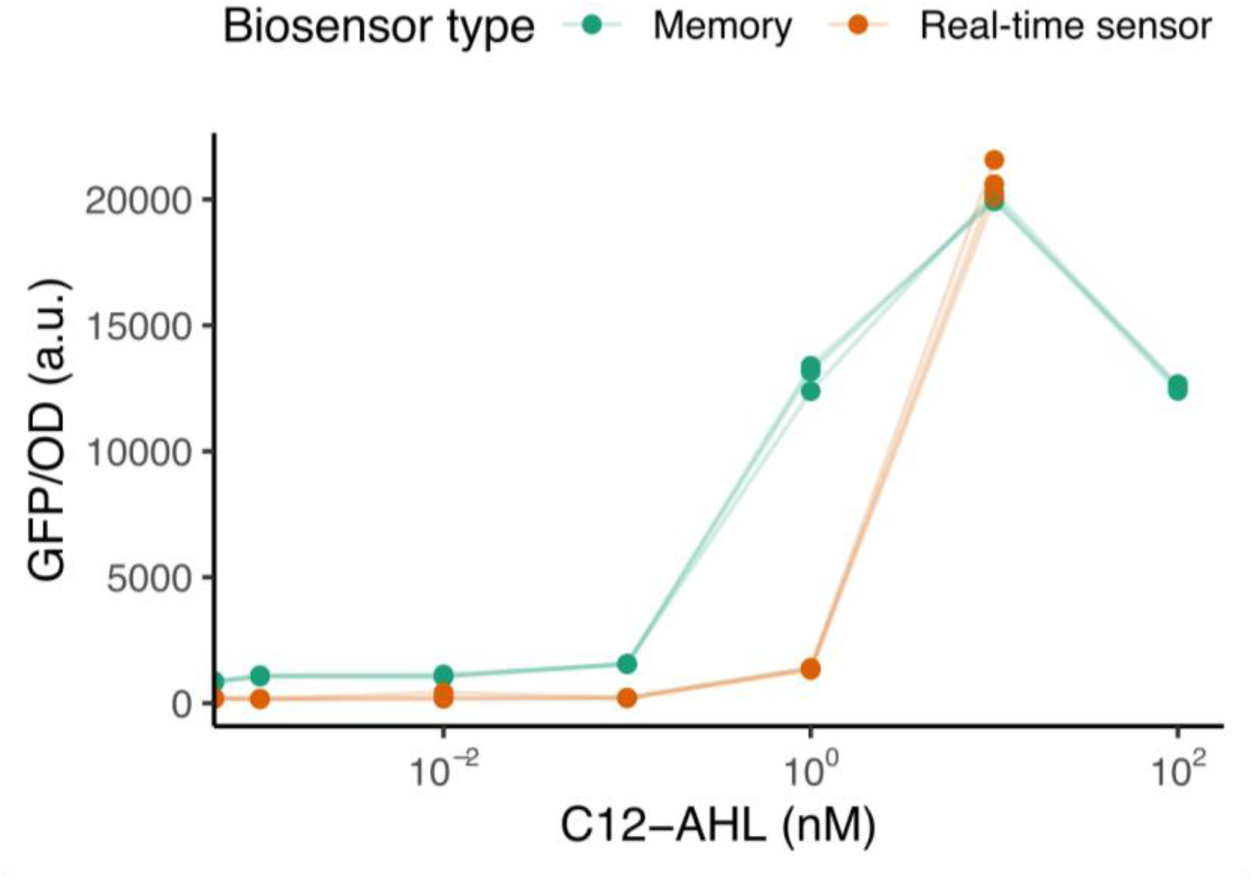
Analog memory response to C12-AHL. We engineered a recombinase-memory sensor for the quorum sensing molecule C12-AHL using the LasR sensor protein and the pLas promoter. The memory-biosensor (green) was tested with different concentrations of C12-AHL to determine sensitivity. The response of a real-time sensor is shown for comparison (orange). The sensor output, shown for three independent replicates, is normalized to cell density (OD600).

**Figure S4.**
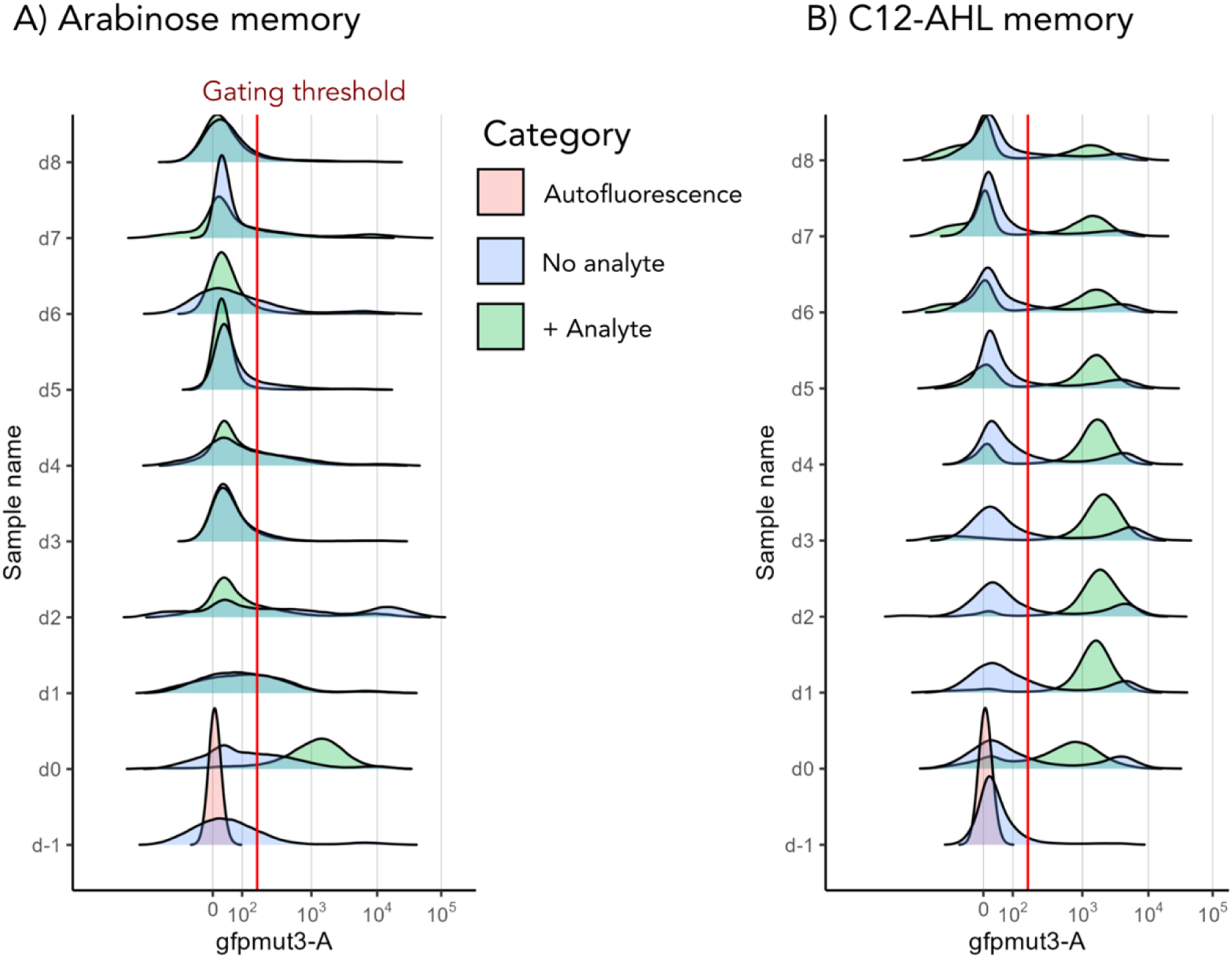
Flow cytometry analysis showing memory loss. Fluorescence distribution showing the ON state populations of (**A**) arabinose and (**B**) C12-AHL memory biosensors over 8 day serial batch cultures. In these ridgeline plots, days are indicated on the y-axis and green fluorescence intensity is shown on the x-axis. Each distribution is a normalized density per sample so the peaks are not comparable across samples. Autofluorescence of MG1655 cells (pink) is shown for reference with the cell populations lacking analyte (blue) and exposed to analyte (green). The threshold for gating (red line) was obtained as 99 percentile of cells with OFF state plasmid only (*data not shown*). We observed that the analyte-exposed populations shifted to higher fluorescence on day 0 (d0) and gradually decreased over time.

**Figure S5.**
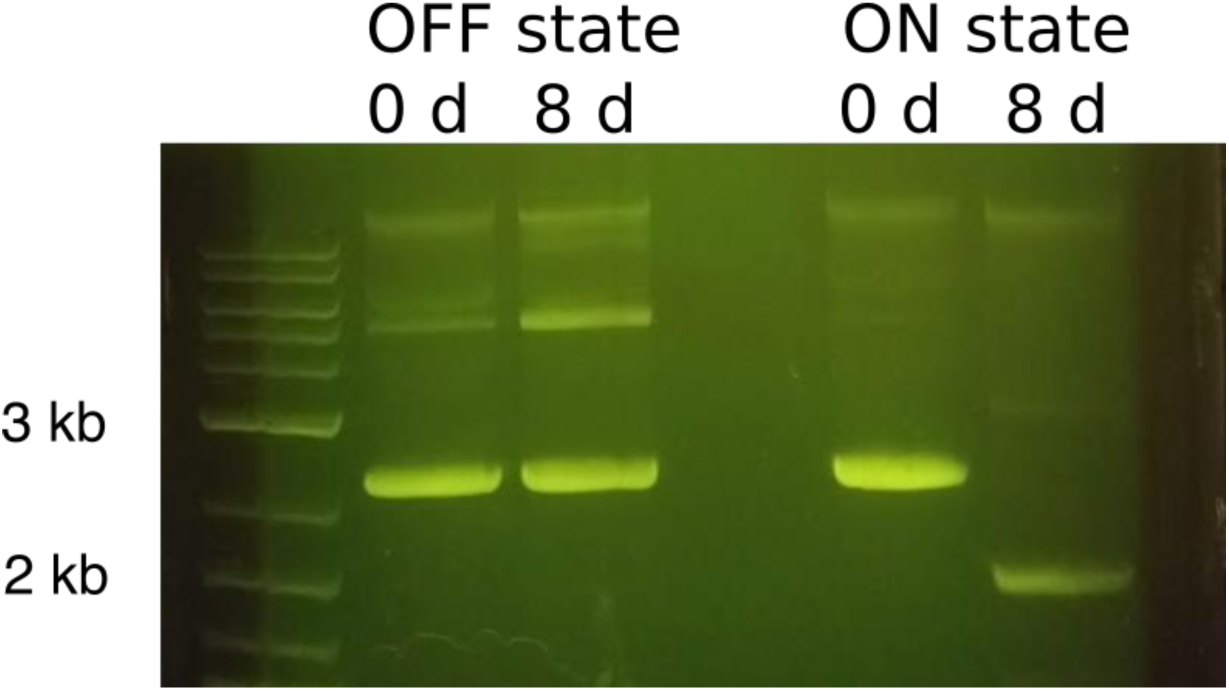
A genetic deletion in the plasmid leads to memory instability. We analyzed the memory-reporter plasmid containing the flipping region to understand if plasmid mutations were responsible for memory loss. We conducted the long duration memory assay on cultures containing only reporter plasmids in both states (pRec1-OFF and pRec1-ON) without the sensor plasmid. We grew 25 ml cultures of the population from day 0 and day 8, extracted the plasmid and ran on agarose gel. We found that the ON state plasmid has a large deletion (∼1.4 kb), which causes the loss of both the fluorescence and the flipping region over the 8 days. We designate this mutated memory plasmid as the BROKEN state.

**Figure S6.**
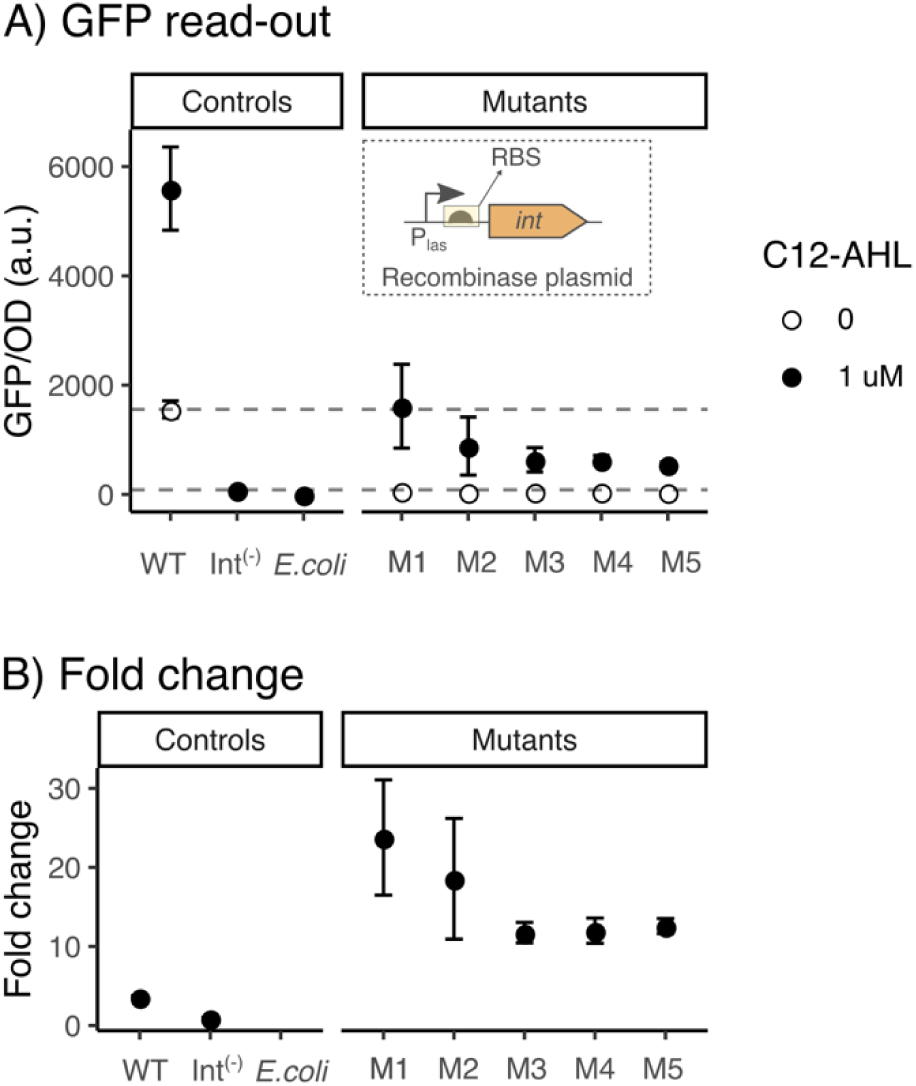
Effect of decreasing recombinase expression on memory performance. (**A**) We created a library of plasmids with altered RBS sequences in the translation initiation sequence used to regulate recombinase expression (inset) and characterized a subset of variants. The plot shows the performance of the 5 memory sensor variants using fluorescence in the presence (closed circles) and absence (open circles) of the analyte C12-AHL. (**B**) Fold change in sensor output with and without the analyte (the ratio of the closed circles to open circles). All of the variants had lower baselines than the starting genetic circuit (WT) in the absence of analyte, a lower maximum normalized fluorescence in the presence of analyte, and a significantly higher fold change (Welch one-tailed t-test, p < 0.04). Data points show the mean and error bars the standard deviation of 3 biological replicates. Int(-) indicates cells lack recombinase.

**Figure S7.**
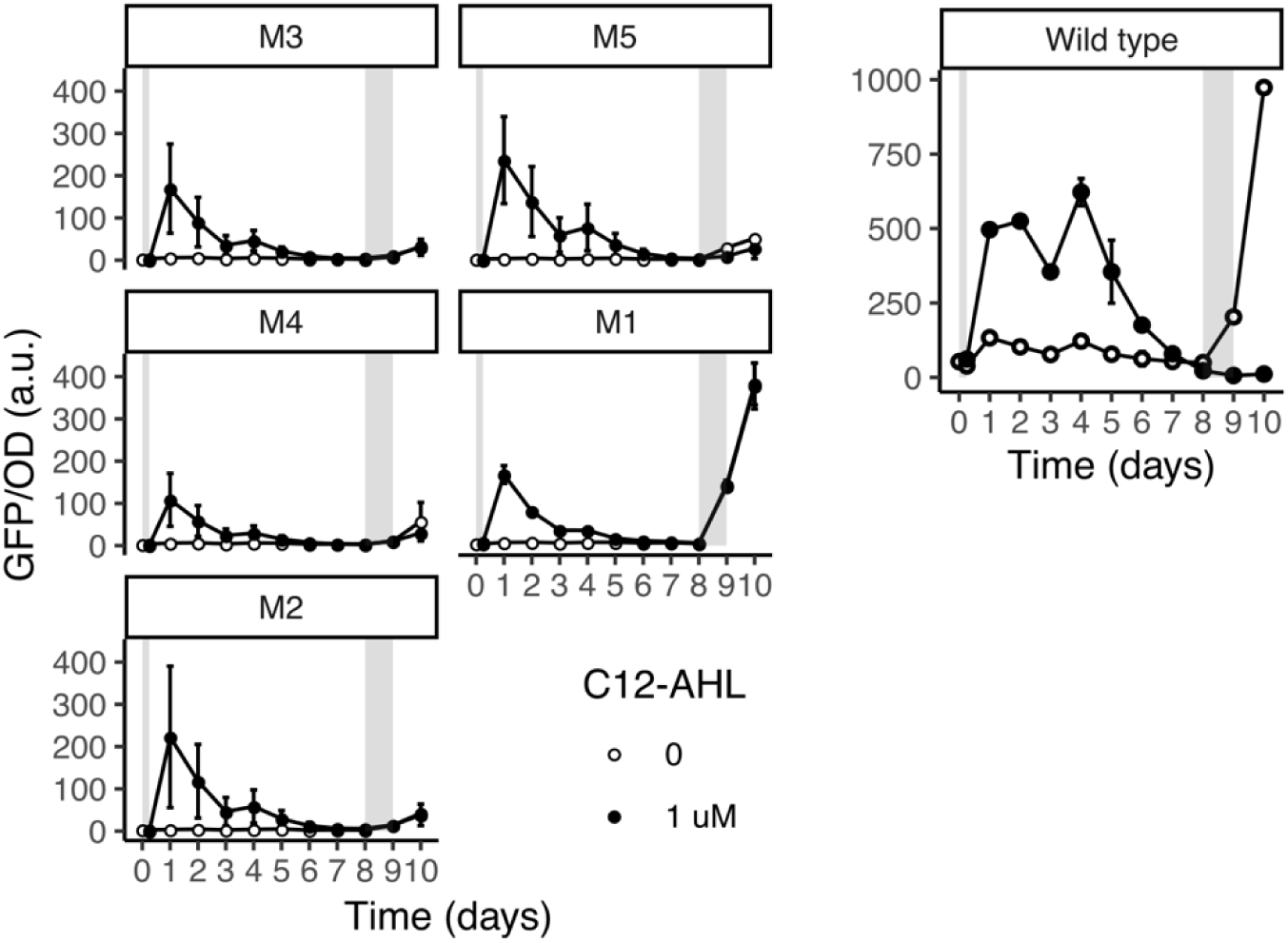
Detection of a second C12-AHL pulse. We tested the performance of RBS variants from Figure S6 (M1 to M5) by adding a second pulse of the analyte C12-AHL on day 8 to both the naive (open circles) and analyte-exposed (closed circles) populations. We compared these results to the original genetic circuit (Wild type). The wild type did not respond to the second pulse if exposed to analyte previously, whereas some mutants responded to the second analyte pulse.

**Figure S8.**
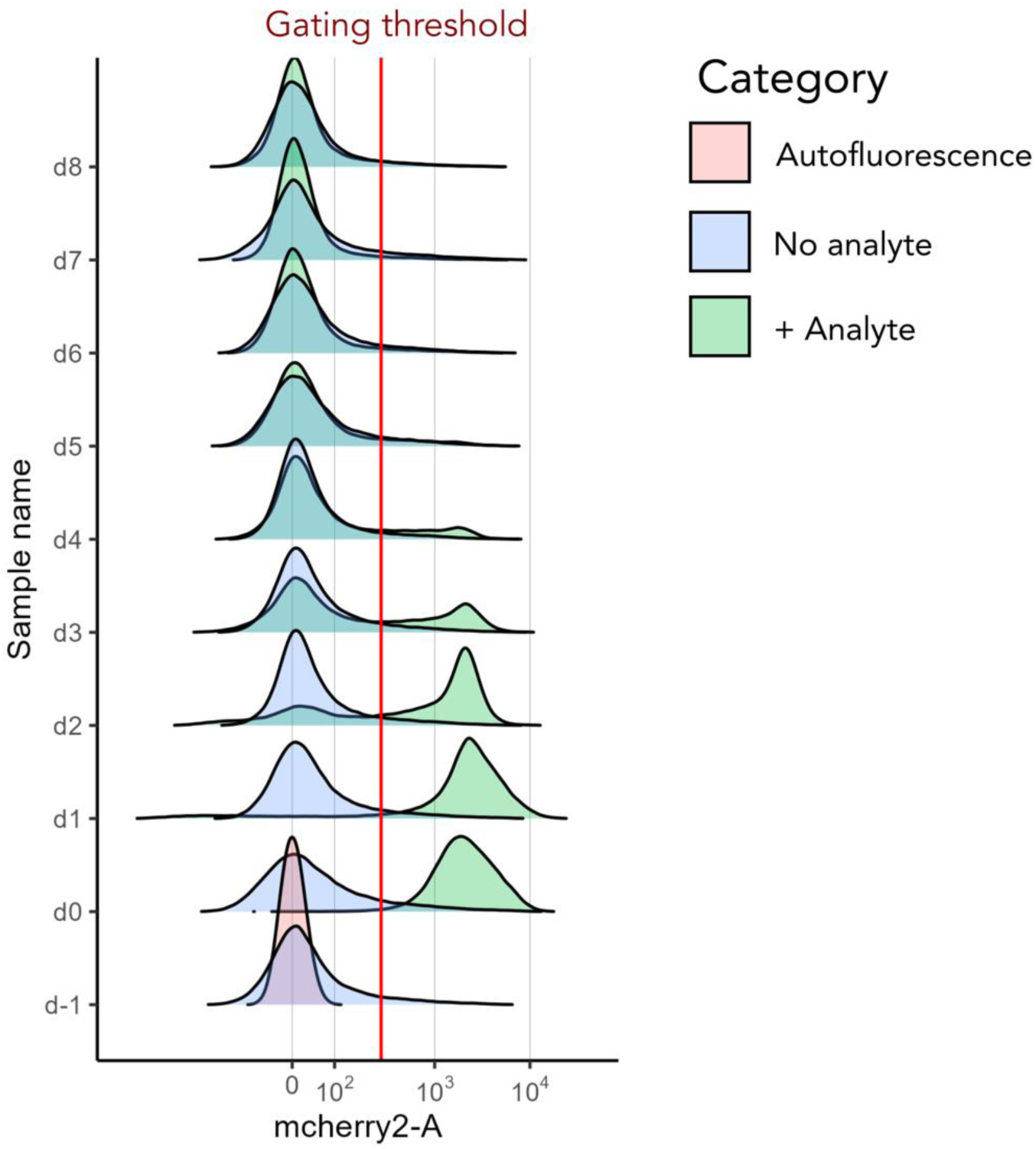
Flow cytometry analysis of the new fluorescent design. Fluorescence distributions showing the ON state populations of the redesigned fluorescent memory biosensor using a red fluorescence output. The ridgeline plots show fluorescence distribution as in Figure S4. Events were gated manually at 300 arbitrary units of fluorescence intensity, such that all the analyte exposed population peaks were above the threshold.

**Figure S9.**
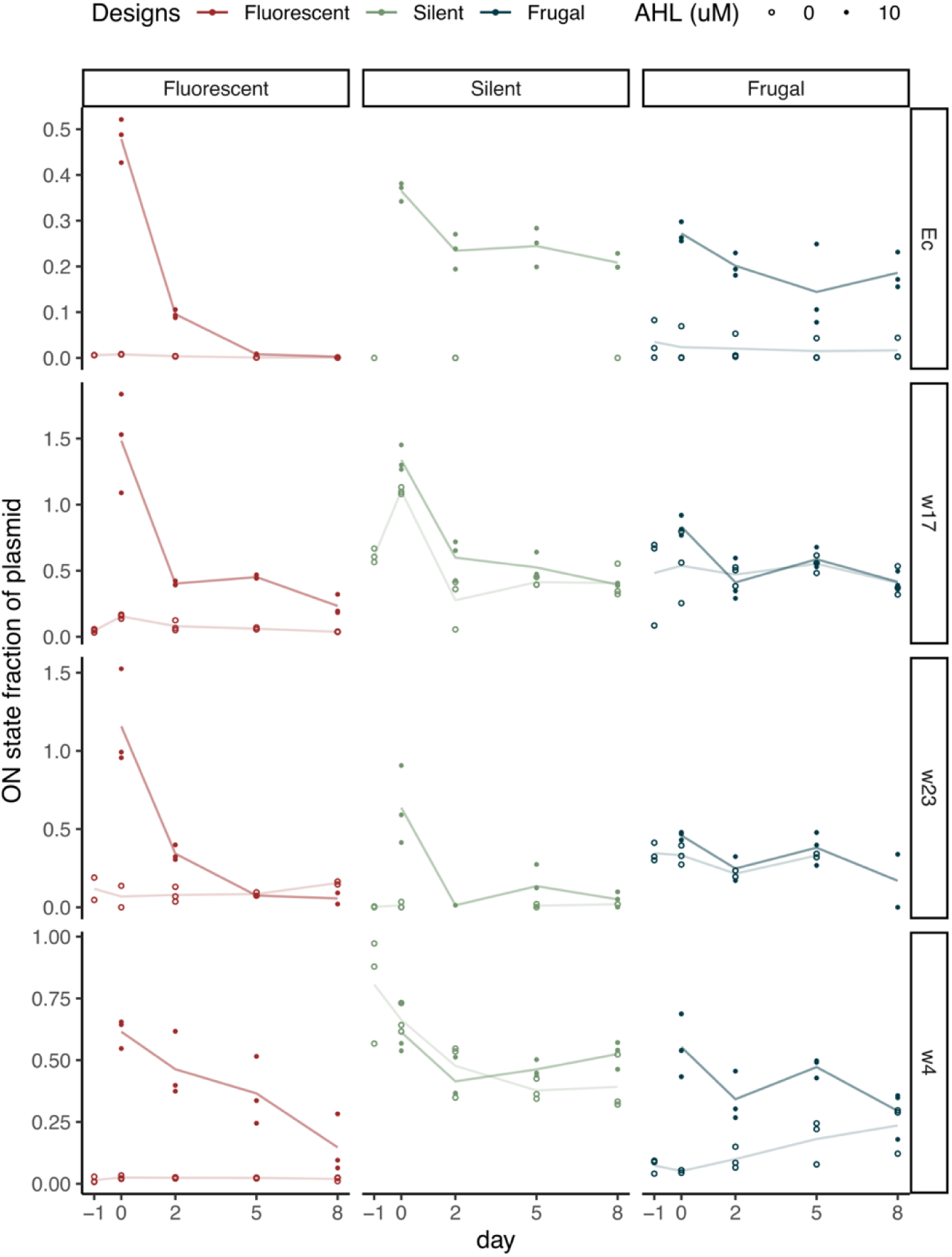
Application of memory in different wastewater isolates. We tested the memory readout over 8 days using qPCR in *E. coli* and three wastewater isolates (w4, w17, w23). The three memory designs are shown in side-by-side panels for each isolate, with the Fluorescent design in red, the Silent design in green, and the Frugal design in blue. The x-axis shows the days of incubation following analyte exposure, while the y-axis shows the fraction of plasmid in the ON state, [copies flipped / total plasmid copies], with data from three independent cultures shown as points, and the line connecting their averages. All organisms have the same memory loss with the fluorescent designs. In the Silent and Frugal designs, *E. coli* and w4 were stable but w17 and w23 had values similar to the samples without exposure to C12-AHL.

**Figure S10.**
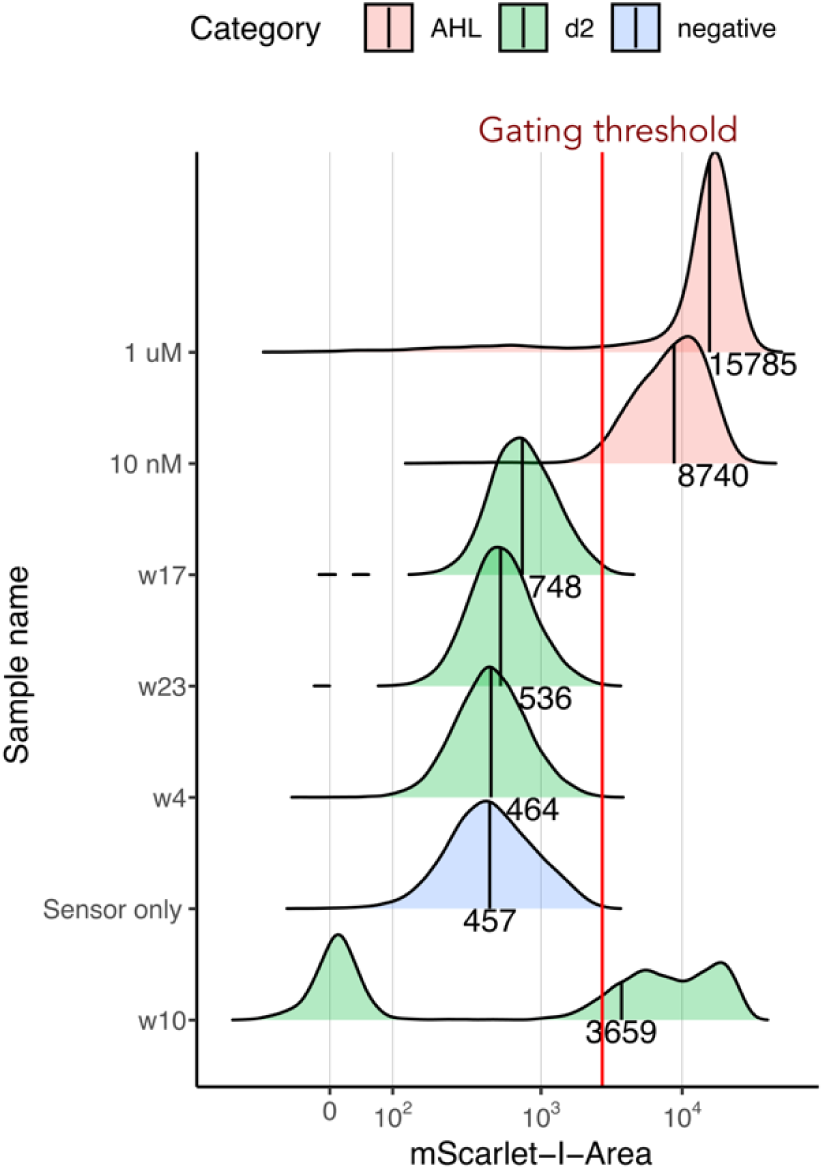
Assaying C12-AHL production by wastewater isolates. We tested for wastewater isolate AHL production by incubating an *E. coli* fluorescent sensor with the spent medium from wastewater isolate cultures. We measured the signal of the E. coli sensor using flow cytometry. Ridgeline plots show the aggregate density distribution of three independent replicate cultures of each sample where fluorescence intensity is on the x-axis and the different samples are distributed across the y-axis. The vertical line within the distribution and the text below indicate the median intensity values for each peak. The red line shows the gating threshold at 99 percentile of the sensor only (negative control) population. Isolates w4, w17 and w23 did not present detectable AHL.

**Figure S11.**
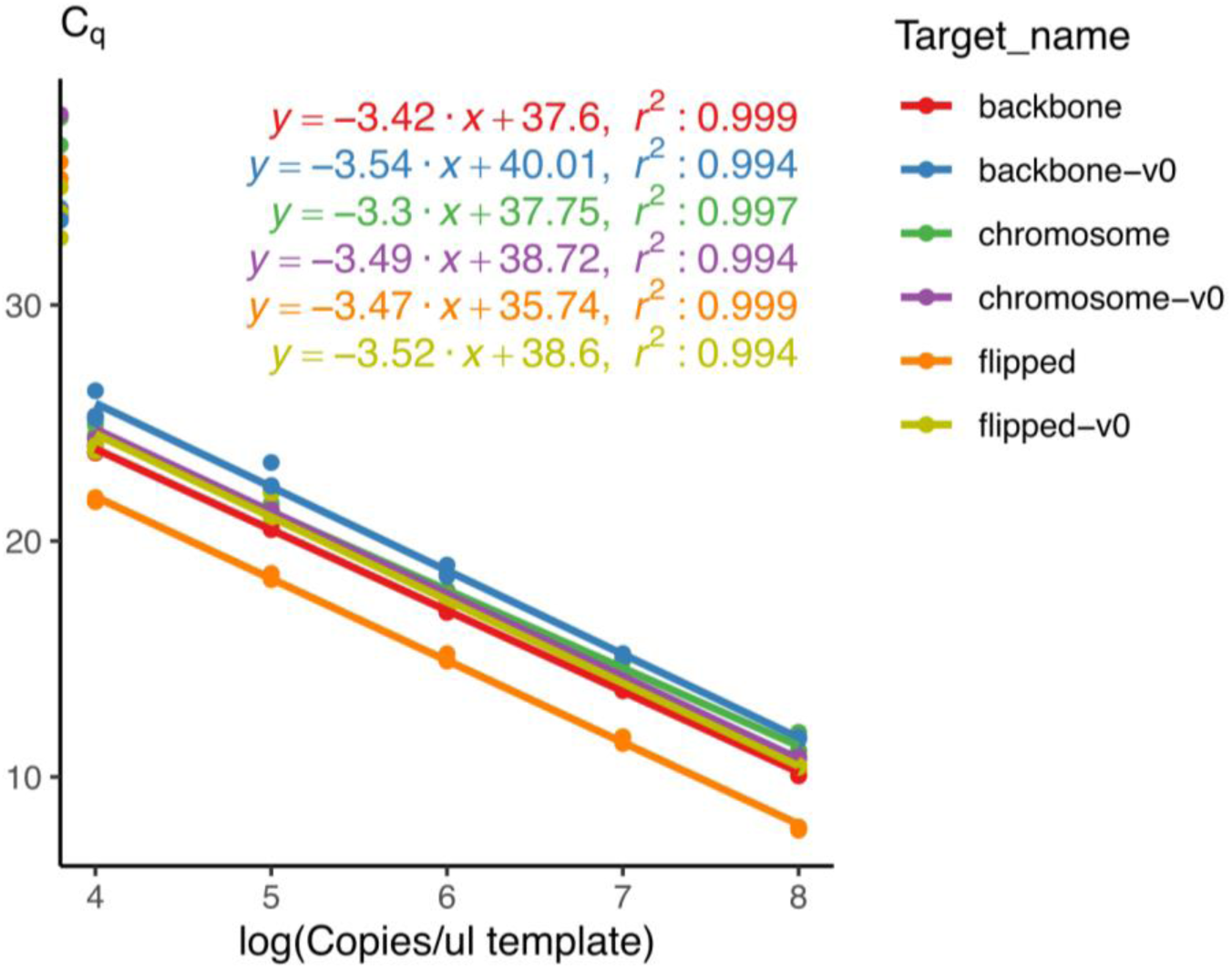
Standard curves for qPCR calibration. We prepared 10-fold serial dilutions of the pooled PCR products of each primer pair in the multiplex reaction to generate standard curves. We gel purified the PCR products, quantified them using a Qubit Broad range kit (Thermo Fisher Scientific), and normalized each of PCR products to achieve a target concentration of 10^8^ copies per µl concentration in the pooled dilution series. We performed multiplex qPCR with the two different sets of 3 targets each, and fitted the Cq values called by the Quantstudio 3 software to standard curve equations shown above the plot. V0 versions target the original designs used in Figure 2D, S2 and other versions target the optimized designs used in Figures 4C, 5B and S9. Amplifications of NTCs where the diluent was run without any target are shown at the left-most end of the plot. PCR efficiencies were between 92% and 101%. Refer to Table S3 for primer sequences.

**Table S1.**
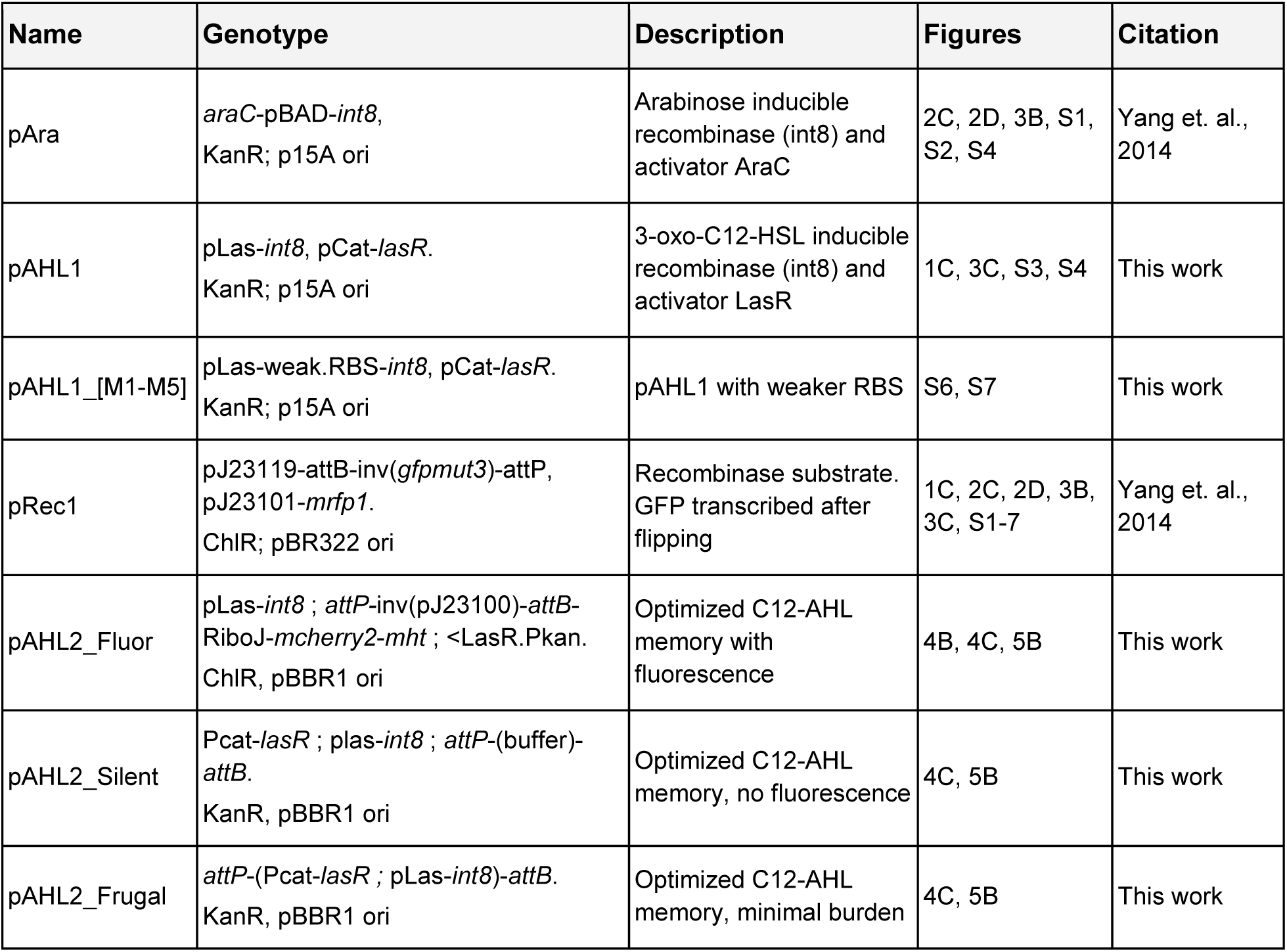
Plasmid list. The table lists the genotype of plasmids used in this work along with a short description and the figures they were used in, along with the citation.

**Table S2.**
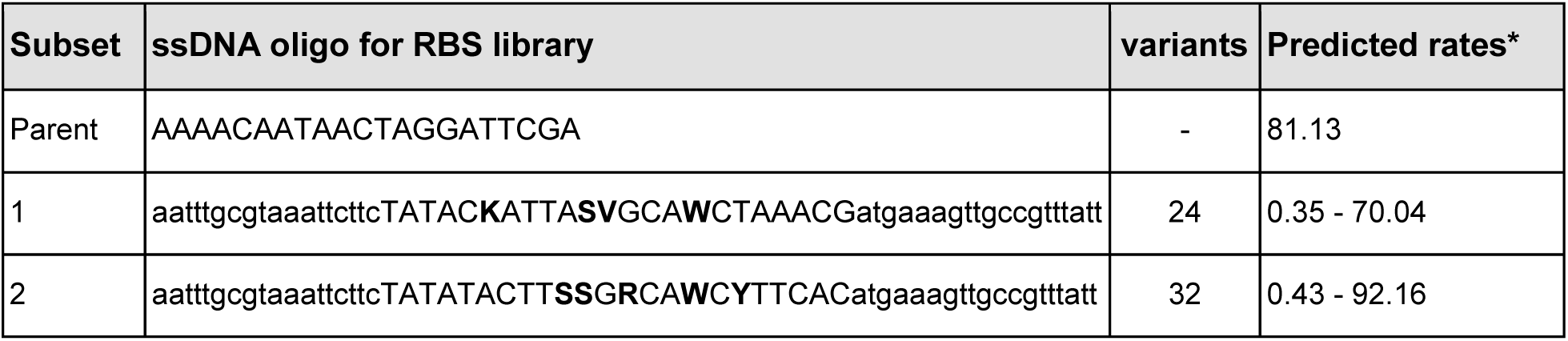
Ribosome binding site variant library. The RBS of the parent is indicated along with the ssDNA oligos used for making the two independent RBS libraries to reduce translation initiation rate of the recombinase (*int8*). The number of variants in each library along with the predicted translation initiation rates (* predicted translation initiation rates from RBS calculator version 2.1). In the ssDNA, 5’ UTR + RBS sequence is capitalized and the flanking regions on both sides are indicated in small letters for reference. Also note the degenerate codons mean the following nucleotides: (K = G/T; S = G/C; V = A/G/C; W = A/T; R = A/G; Y = C/T).

**Table S3.**
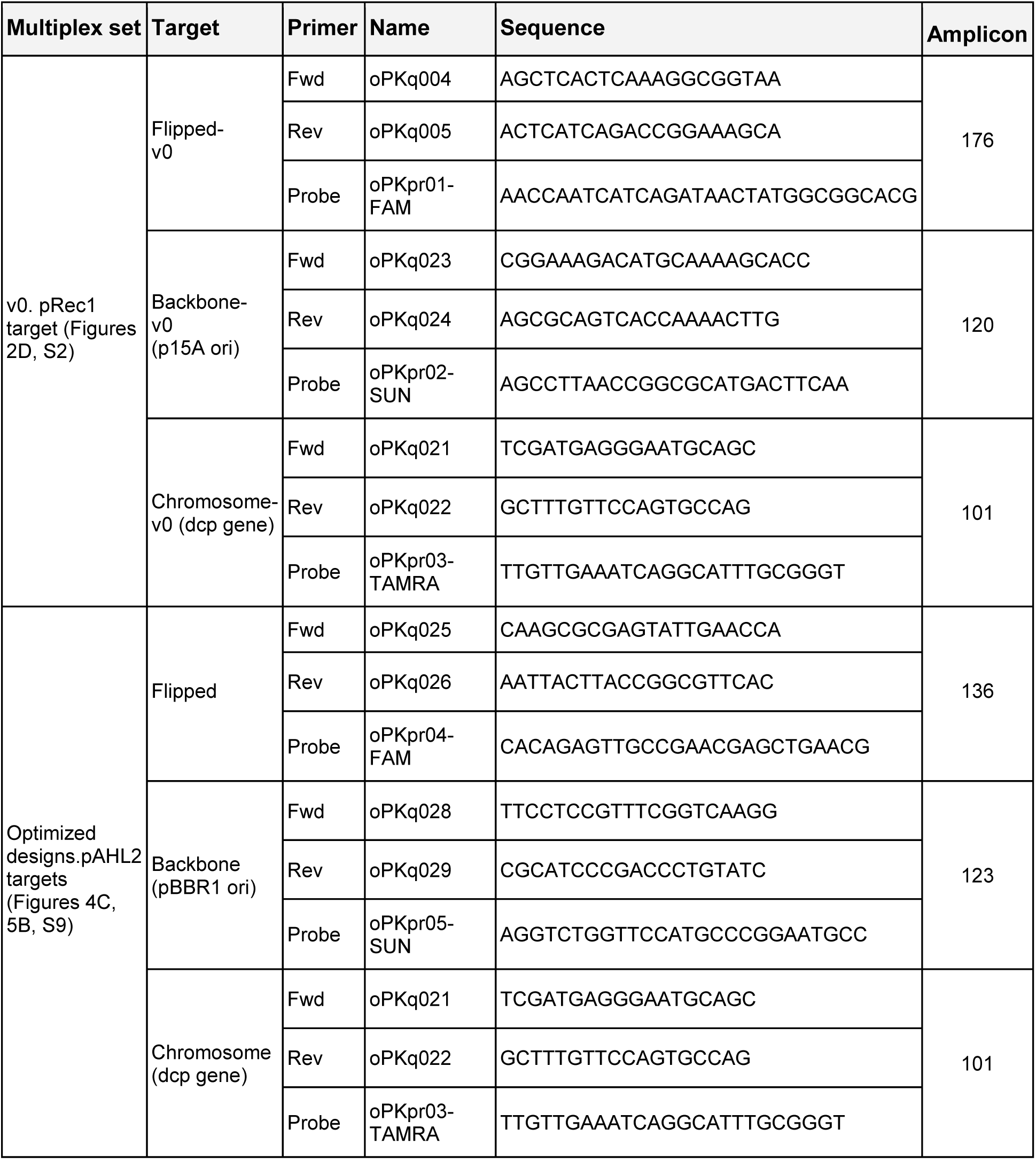
Primer list for qPCR. The table lists each triplex reaction set, with three targets along with the figures they were used in. The forward and reverse primers, and the hydrolysis probe for each target are indicated by name and DNA sequence. Probes included a 5’ fluorophore, FAM and SUN probes included an internal ZEN quencher at 9th bp, and a 3’ Iowa Black FQ quencher, and TAMRA probes included a 3’ BHQ-1 quencher. *E. coli dcp* gene (Uniprot : P24171) was chosen to measure chromosomal copy number, due to close proximity to replication terminus terC, expected to be nearly one per cell in any growth phase (1,2).

**Table S4.**
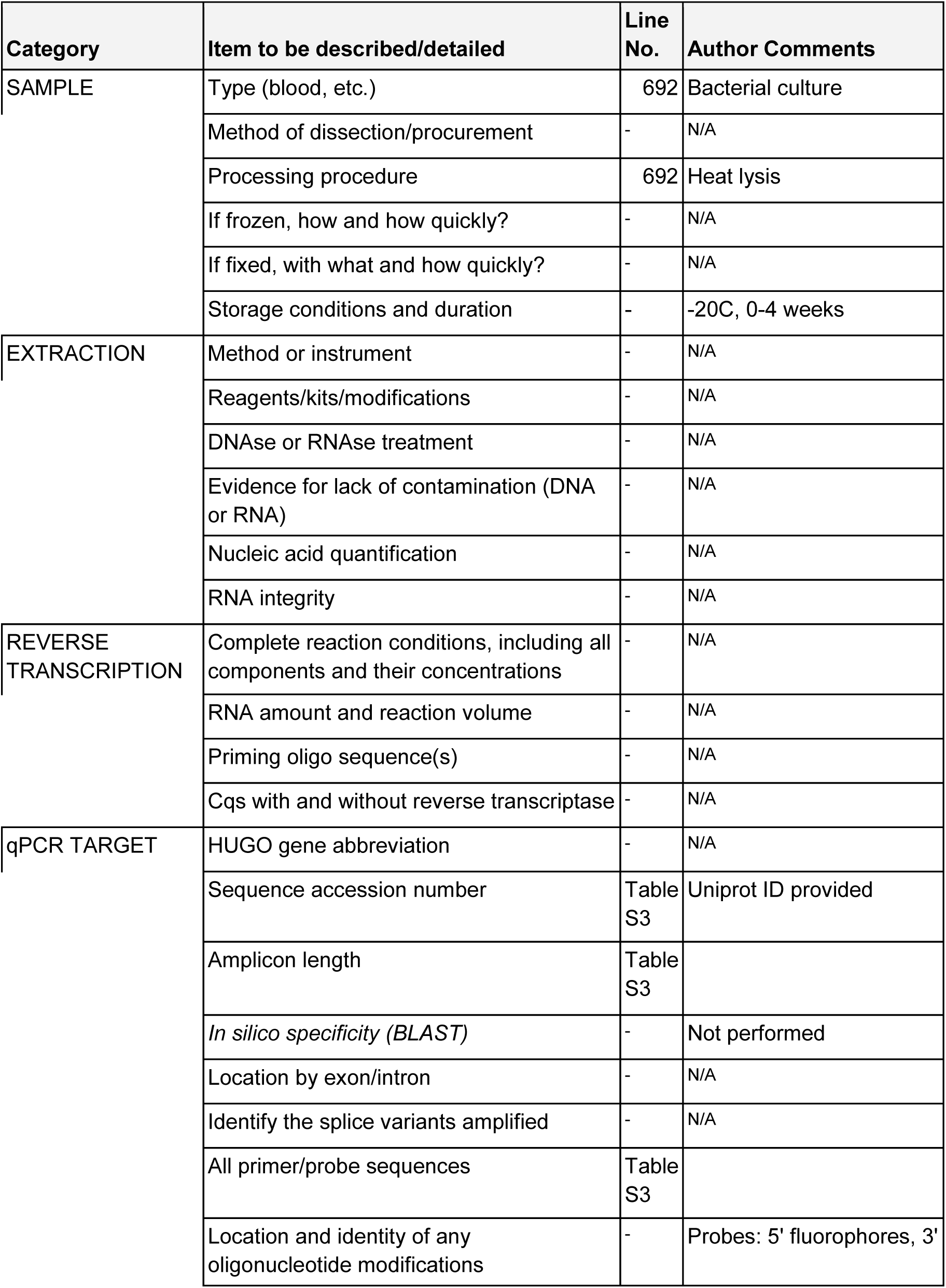
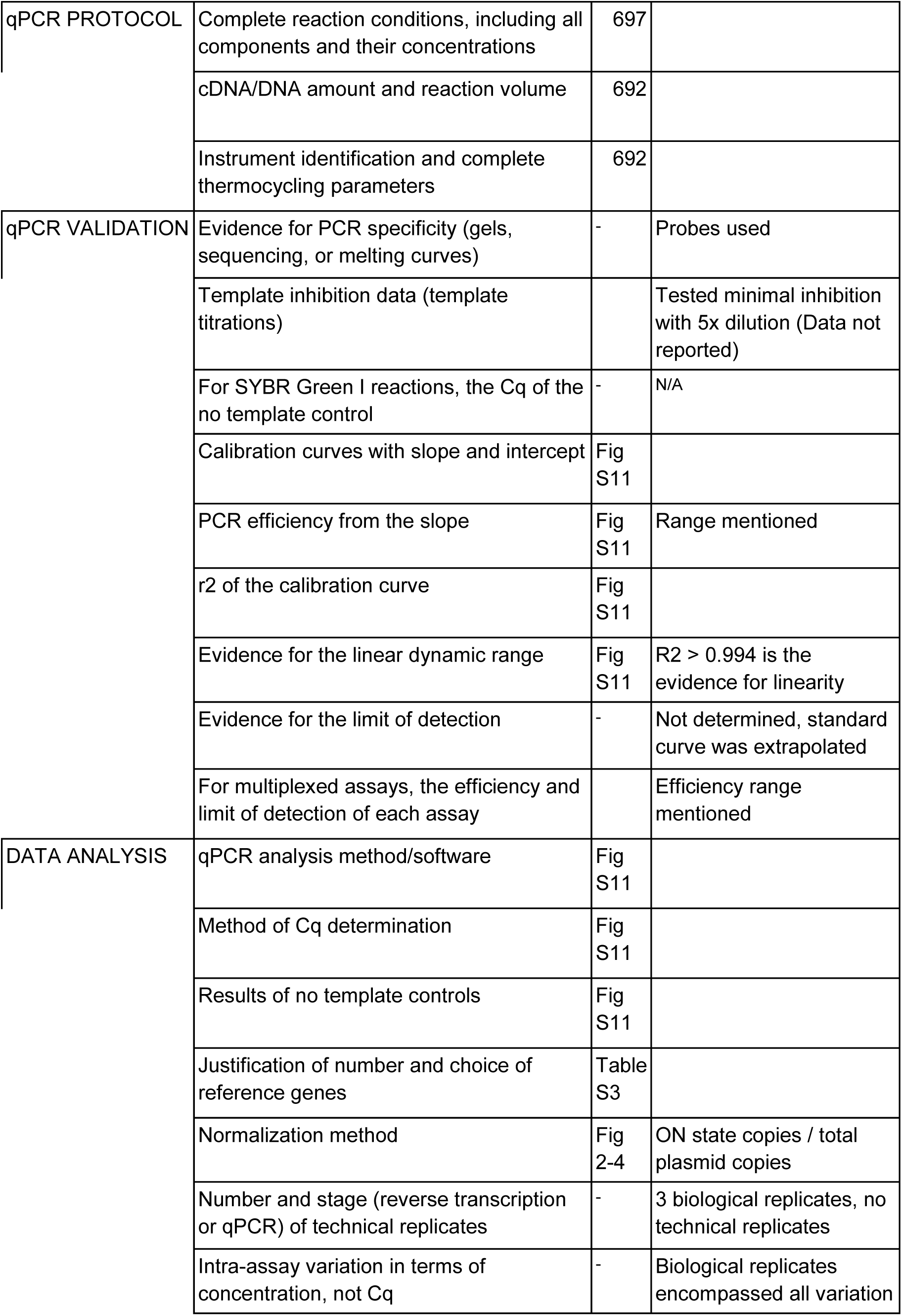
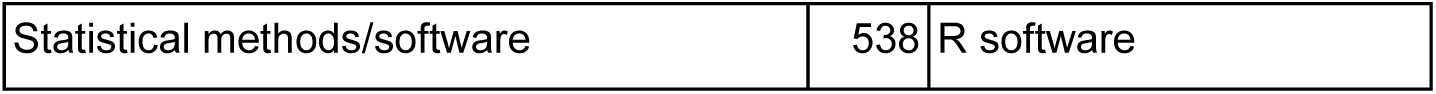
MIQE checklist for qPCR. Checklist summarizing qPCR procedures, sample processing and data processing mentioned within the manuscript, in accordance with the MIQE guidelines, *i.e.*, minimum information for publication of qPCR experiments (3).

| Category | Item to be described/detailed | Line No. | Author Comments |
| --- | --- | --- | --- |
| SAMPLE | Type (blood, etc.) | 692 | Bacterial culture |
|  | Method of dissection/procurement | - | N/A |
|  | Processing procedure | 692 | Heat lysis |
|  | If frozen, how and how quickly? | - | N/A |
|  | If fixed, with what and how quickly? | - | N/A |
|  | Storage conditions and duration | - | -20C, 0-4 weeks |
| EXTRACTION | Method or instrument | - | N/A |
|  | Reagents/kits/modifications | - | N/A |
|  | DNAse or RNAse treatment | - | N/A |
|  | Evidence for lack of contamination (DNA or RNA) | - | N/A |
|  | Nucleic acid quantification | - | N/A |
|  | RNA integrity | - | N/A |
| REVERSE TRANSCRIPTION | Complete reaction conditions, including all components and their concentrations | - | N/A |
|  | RNA amount and reaction volume | - | N/A |
|  | Priming oligo sequence(s) | - | N/A |
|  | Cqs with and without reverse transcriptase | - | N/A |
| qPCR TARGET | HUGO gene abbreviation | - | N/A |
|  | Sequence accession number | Table S3 | Uniprot ID provided |
|  | Amplicon length | Table S3 |  |
|  | *In silico* specificity (BLAST) | - | Not performed |
|  | Location by exon/intron | - | N/A |
|  | Identify the splice variants amplified | - | N/A |
|  | All primer/probe sequences | Table S3 |  |
|  | Location and identity of any oligonucleotide modifications | - | Probes: 5' fluorophores, 3' |
| qPCR PROTOCOL | Complete reaction conditions, including all components and their concentrations | 697 |  |
|  | cDNA/DNA amount and reaction volume | 692 |  |
|  | Instrument identification and complete thermocycling parameters | 692 |  |
| qPCR VALIDATION | Evidence for PCR specificity (gels, sequencing, or melting curves) | - | Probes used |
|  | Template inhibition data (template titrations) |  | Tested minimal inhibition with 5x dilution (Data not reported) |
|  | For SYBR Green I reactions, the C<sub>q</sub> of the no template control | - | N/A |
|  | Calibration curves with slope and intercept | Fig S11 |  |
|  | PCR efficiency from the slope | Fig S11 | Range mentioned |
|  | r<sup>2</sup> of the calibration curve | Fig S11 |  |
|  | Evidence for the linear dynamic range | Fig S11 | R<sup>2</sup> > 0.994 is the evidence for linearity |
|  | Evidence for the limit of detection | - | Not determined, standard curve was extrapolated |
|  | For multiplexed assays, the efficiency and limit of detection of each assay |  | Efficiency range mentioned |
| DATA ANALYSIS | qPCR analysis method/software | Fig S11 |  |
|  | Method of C<sub>q</sub> determination | Fig S11 |  |
|  | Results of no template controls | Fig S11 |  |
|  | Justification of number and choice of reference genes | Table S3 |  |
|  | Normalization method | Fig 2-4 | ON state copies / total plasmid copies |
|  | Number and stage (reverse transcription or qPCR) of technical replicates | - | 3 biological replicates, no technical replicates |
|  | Intra-assay variation in terms of concentration, not C<sub>q</sub> | - | Biological replicates encompassed all variation |

|  |  |  |
| --- | --- | --- |
| Statistical methods/software | 538 | R software |

